# Hippocampal mGluR5 levels are comparable in Alzheimer’s and control brains, and divergently influenced by amyloid and tau in control brain

**DOI:** 10.1101/2024.05.25.595868

**Authors:** Junlong Wang, Serena Savodalli, Yanyan Kong, Cinzia A. Maschio, Uwe Konietzko, Jan Klohs, Daniel Razansky, Axel Rominger, Linjing Mu, Roger Schibli, Christoph Hock, Roger M. Nitsch, Ruiqing Ni

## Abstract

**Background:** Metabotropic glutamate receptor 5 (mGluR5) modulates excitatory glutamatergic synaptic transmission and plays an important role in learning and memory formation and in neurodegeneration and amyloid deposition in Alzheimer’s disease (AD). Conflicting results on the cerebral mGluR5 levels in AD have been reported based on *in vivo* and postmortem studies. Here, we aimed to assess alterations in hippocampal mGluR5 expression in AD, and the associations between mGluR5 expression and pathologies.

**Methods:** Immunofluorescence staining for mGluR5 was performed on postmortem brain tissue from 34 AD patients and 31 nondemented controls (NCs) and from aged 3×Tg and arcAβ model mice of AD. Autoradiography was performed on brain tissue slices from arcAβ mice using mGluR5 tracer [^18^F]PSS232. Analysis of different cellular source of GRM5 RNA in human and mouse brains was performed. Proteomic profiling and pathway analysis were performed on hippocampal tissue from aged 3×Tg mice and wild-type mice.

**Results:** No differences in hippocampal mGluR5 expression or entorhinal cortical GRM5 RNA levels were detected between the AD and NC groups. Hippocampal mGluR5 levels increased with Braak stage and decreased with amyloid level in the NC group. No correlations were detected between the levels of mGluR5 and amyloid, tau, or Iba1/P2X7R in the hippocampus of AD patients and NC cases. *Ex vivo* autoradiography revealed comparable cerebral levels of [^18^F]PSS232 in arcAβ mice compared to nontransgenic mice. GO and KEGG pathway enrichment analyses revealed that the Shank3, Grm5 and glutamatergic pathways were upregulated in hippocampal tissue from aged 3×Tg mice compared to wild-type mice.

**Conclusion:** This study revealed no difference in hippocampal mGluR5 levels between AD patients and NCs and revealed the divergent influence of amyloid and tau pathologies on hippocampal mGluR5 levels in NCs. Species differences were observed in the GRM5 RNA level as well as at the cellular location.

**Graphical abstract:** 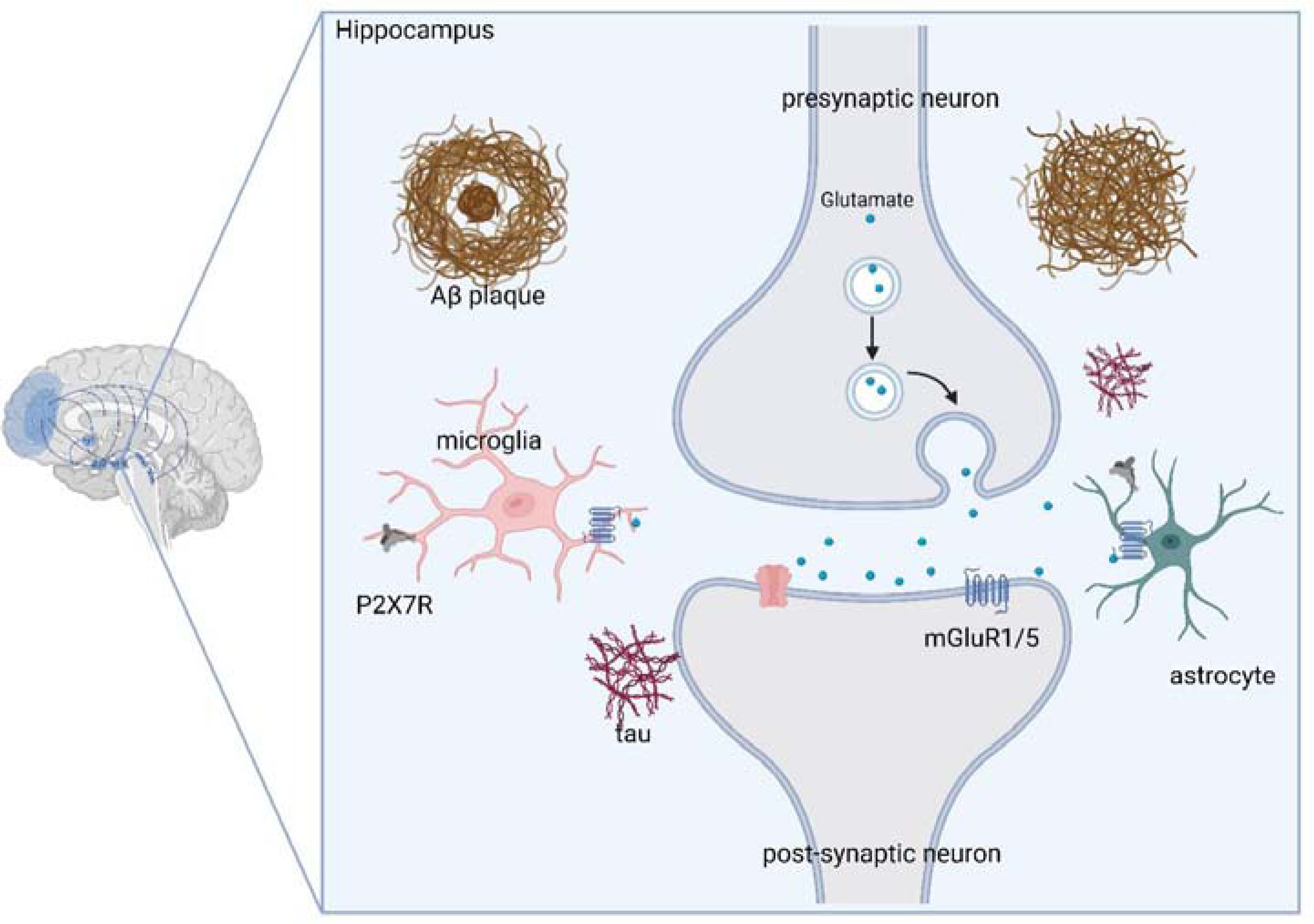

## Introduction

Alzheimer’s disease (AD) is pathologically characterized by the aberrant accumulation of amyloid-beta (Aβ), tau tangles, neuroinflammation, and synaptic dysfunctions (9, 32, 35). Synaptic degeneration and neurotransmitter receptor alterations play critical roles in the progression of AD (54, 72). Glutamatergic neurons form the main excitatory system in the brain and play a pivotal role in many neurophysiological functions. Excessive activation of glutamate receptors can lead to excitotoxicity, neuronal dysfunction and death, impaired calcium buffering, free radical generation, and mitochondrial permeability transition in the brain (64). Metabotropic glutamate receptor 5 (mGluR5) is a G protein-coupled receptor located predominantly at the postsynaptic terminal and in the presynaptic membrane, as well as on microglia (24, 48). In healthy adult astrocytes, mGluR5 is generally absent but transiently reemerges in astrocytes within a restricted time frame. In the brain, mGluR5 is mainly expressed in the hippocampus, amygdala, olfactory bulb, dorsal striatum, nucleus accumbens, and lateral septum (67). mGluR5 modulates excitatory glutamatergic synaptic transmission and plays an important role in learning, memory and neuronal development and glutamate-induced excitotoxicity (24, 48). mGluR5 has implications for the pathophysiology of many neurodegenerative diseases (10), including AD (2, 46). Aβ oligomers have been shown to be toxic and can lead to synaptic impairment (68). mGluR5 has been shown to act as a coreceptor for Aβ oligomers that bind to cellular prion proteins (57, 73), mediating synaptic dysfunction (57). Soluble Aβ oligomers have been shown to induce the accumulation and overactivation of mGluR5, leading to an abnormal increase in the release of intracellular Ca2+ (57, 73). Genetic deletion of mGluR5 has been shown to improve cognitive function and reduce Aβ plaque, Aβ oligomer accumulation, and mTOR phosphorylation in the brains of APP/PS1 mice of AD (22). Pharmacological treatment, such as allosteric modulation of mGluR5, prevents cognitive impairment, reduces pathogenesis, and reverses synapse loss in AppNL-G-F/hMapt double-knock-in mice and APP/PS1 mouse models of AD (21, 23, 65). Chronic pharmacological inhibition of mGluR5 has also been shown to prevent cognitive impairment and reduce pathogenesis in AD mice (23). Moreover, mGluR5 has been implicated in modulating microglial inflammation, such as in Parkinson’s disease (PD) (84).

Given its important pathophysiological role, mGluR5 is thus a promising target in drug development and a promising imaging biomarker for neurodegenerative diseases, including AD (53). Several mGluR5 positron emission tomography (PET) tracers, including [^18^F]FPEB (80), [^11^C]ABP688 (8) and [^18^F]PSS232 (60, 78), have been developed. Silent allosteric modulator (SAM) and negative allosteric modulators (NAMs) of mGluR5 are currently being tested in clinical trials in AD patients as well as in PD patients (e.g., NCT05804383; NCT04857359). However, existing results from *in vivo* and *ex vivo* studies on changes in mGluR5 expression in the AD brain are conflicting. By western blotting, a significant decrease in the protein expression of mGluR1, but not mGluR5, was detected in the frontal cortex tissue of AD patients (7). A phosphoproteomic study revealed that the GRM-calcium signaling pathway in the frontal cortex was more enriched in patients with AD than in controls (59). Autoradiography revealed increased [^18^F]PSS232 binding in the cortex and hippocampus of 6 AD patients compared to 6 NC subjects (46). However, another autoradiography study using L- [^3^H]glutamate revealed a reduced binding level in the frontal cortex of AD patients, which directly correlated with Braak stage progression in AD patients (7). Thus far, four *in vivo* studies have been conducted, and reduced uptake was shown in the hippocampus of AD patients compared to that in NC subjects in relatively small cohorts by using [^11^C]ABP688 (70), [^18^F]FPEB (44), and [^18^F]PSS232 (76, 77). In addition, little is known about the stage dependency and influence of APOE on mGluR5 levels in the AD brain and its link with tau deposits in the brain. Moreover, mGluR5 is one of the major mGluRs identified on microglia and astrocytes (84). The activation of microglial mGluR5 has been shown to regulate neuroinflammation in disease animal models (52, 75). However, the association between the level of mGluR5 and microglial activation in the AD brain has not been directly quantified.

Given the conflicting results, the aim of the current study was to assess alterations in hippocampal mGluR5 expression and RNA levels in AD patients and NC subjects and in tissue from transgenic animal models of AD. Furthermore, we aimed to determine the influence of age, sex, and the APOE e4 allele and to determine the associations of mGluR5 levels with Aβ and tau pathologies and neuroinflammation.

## Materials and Methods

### Animal models

Six 3×Tg mice [B6;129-Psen1tm1MpmTg(APPSwe, tauP301L)1Lfa/Mmjax] were purchased from Jax Laboratory (33, 36). Ten arcAβ transgenic mice (human APP695 transgenes harboring Swedish and Arctic (E693G) mutations under the control of the prion protein promoter) (30, 56) and eight nontransgenic littermates were used. For the proteomics study, wild-type C57BL6 mice were obtained from Cavins Laboratory Animal Co., Ltd., of Changzhou. Mice were housed in ventilated cages inside a temperature-controlled room under a 12 h dark/light cycle. Pelleted food and water were provided ad libitum. Paper tissue and red mouse house shelters were placed in cages for environmental enrichment. For the proteomics study, the experimental protocol used was approved by the Institutional Animal Care and Ethics Committee of Huashan Hospital of Fudan University and performed in accordance with the National Research Council’s Guide for the Care and Use of Laboratory Animals. All experiments were carried out in compliance with national laws for animal experimentation and were approved by the Animal Ethics Committee of Fudan University. For the staining and autoradiography studies, the experiments were performed in accordance with the Swiss Federal Act on Animal Protection and were approved by the Cantonal Veterinary Office Zurich. The animals were housed in ventilated cages inside a temperature-controlled room under a 12-h dark/light cycle. Pelleted food (3437PXL15, CARGILL) and water were provided ad libitum. Paper tissue and red Tecniplast Mouse House® (Tecniplast, Milan, Italy) shelters were placed in cages for environmental enrichment. 3×Tg mice, C57B6 wild-type mice, arcAβ mice and nontransgenic mice were perfused under ketamine/xylazine/acepromazine maleate anesthesia (75/10/2 mg/kg body weight, i.p. bolus injection) with ice-cold 0.1 M PBS (pH 7.4) and 4% paraformaldehyde in 0.1 M PBS (pH 7.4), fixed for 24 h in 4% paraformaldehyde (pH 7.4) and then stored in 0.1 M PBS (pH 7.4) at 4°C.

### Postmortem human brain tissue

Postmortem hippocampal brain tissue from thirty-four patients with ADs, each with a clinical diagnosis confirmed by pathological examination, and thirty NC subjects were included in this study (**Table 1**). Autopsy paraffin-embedded hippocampal tissues were obtained from the Netherlands Brain Bank (NBB), Netherlands. All materials were collected from donors or from whom written informed consent was obtained for a brain autopsy, and the use of the materials and clinical information for research purposes were obtained by the NBB. The study was conducted according to the principles of the Declaration of Helsinki and subsequent revisions. All the autopsied human brain tissue experiments were carried out in accordance with ethical permission obtained from the regional human ethics committee and the medical ethics committee of the VU Medical Center for NBB tissue. Information on the neuropathological diagnosis of AD (possible, probable, or definite AD) or not AD was obtained from the NBB. Information on the Consortium to Establish a Registry for AD (CERAD), which applied semiquantitative estimates of neuritic plaque density and the Braak score (9) based on the presence of neurofibrillary tangles (NFTs), is provided in Table 1. Patients with pathology other than AD pathology were excluded from the study.

**Table 1.**
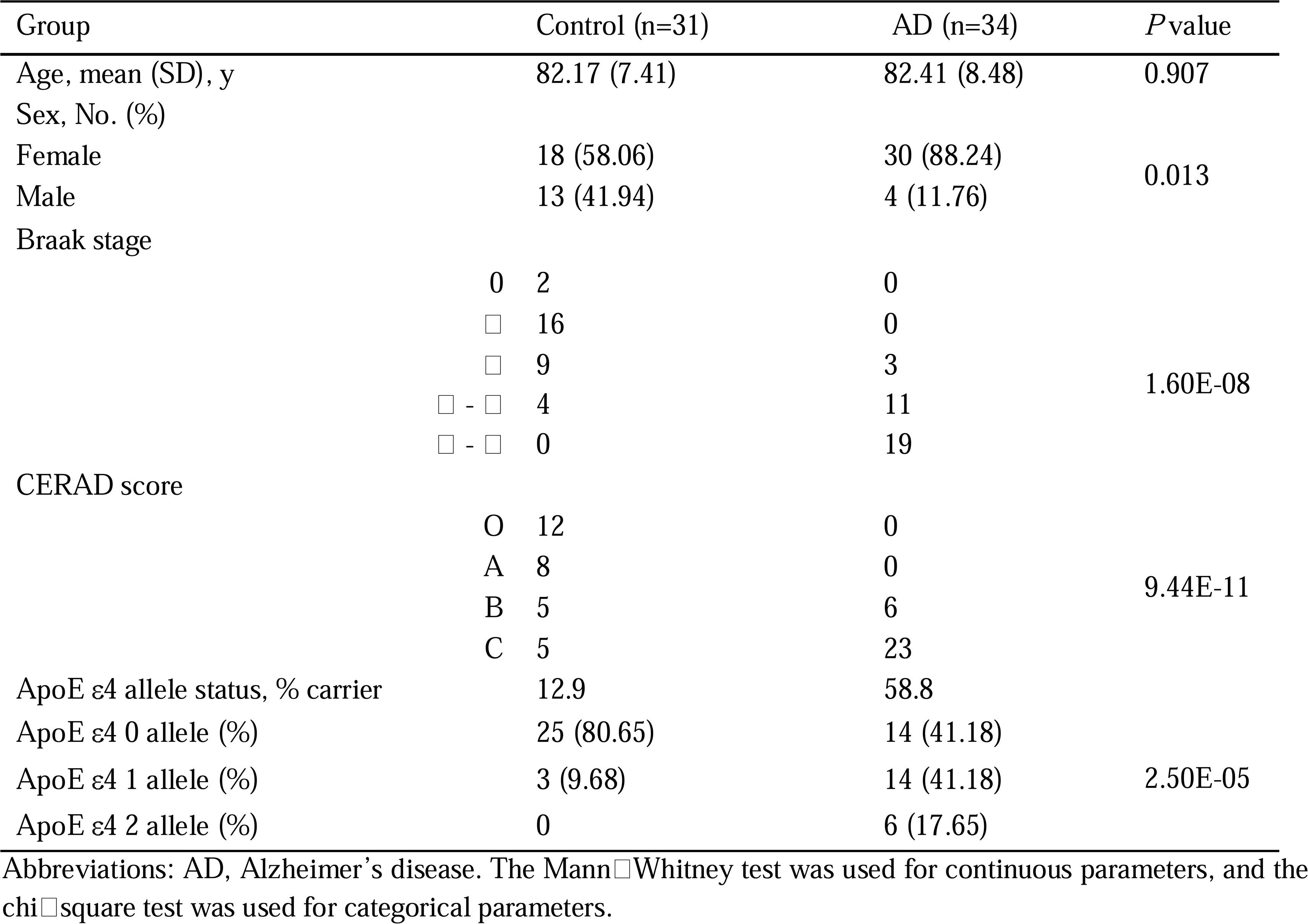
Demographics of postmortem human brains.

### Radiosynthesis

For the mGluR5 tracer [^18^F]PSS232, the molar activity ranged from 46.4 to 73.9 GBq/μmol at the end of synthesis, with a radiochemical purity > 95% (46). In a typical experiment, a moderate radiochemical yield of ∼ 12% (decay corrected) was achieved with a radiochemical purity > 99%. In postsynaptic elements, mGlu5 receptors are physically linked to the NR2 subunit of the N-methyl-D-aspartate receptor (NMDAR) via a chain of interacting proteins, including PSD-95, Shank and Homer (71). The GluN2B subunit is a major NR2 subtype of the NMDA receptor and is highly expressed only in the cortex and hippocampus, with a rather low density in the cerebellum and hypothalamus (51). Here, we assessed the level of GluN2B using [^11^C]WMS1405B (69). The GluN2B subunit is one of the major NR2 subtypes of NMDA receptors and is highly expressed only in the cortex and hippocampus; moreover, the GluN2B subunit is expressed at a rather low density in the cerebellum and hypothalamus (51) and mediates the acute and chronic synaptotoxic effects of oligomeric Aβ in murine models of AD (55). Several PET ligands for GluN2B have been developed, including (S)-[^18^F]OF-NB1 (5), (R)- [^18^F]OF-Me-NB1 (5), (R)-[^11^C]NR2B-Me (13), [^11^C]Me-NB1 (5, 37), [^11^C]Ro04-5595 (26) and [^11^C]WMS-1405B (69). The molar activities of [^11^C]WMS1405B ranged from 53.8 to 295.3 GBq/μmol at the end of synthesis, with a radiochemical purity > 99% (69). The molar activities ranged from 156 to 194 GBq/μmol at the end of synthesis. The identity of the final product was confirmed by comparison with the high-performance liquid chromatography (HPLC) retention time of the nonradioactive reference compound after coinjection (47).

### Autoradiography of postmortem brain tissues from arcA**β** mice

Four arcAβ mice and three NTL mice were used in the autoradiography. Dissected mouse brains embedded in TissueTek were cut into 10-μm-thick sagittal sections on a cryostat (Cryo-Star HM 560 MV; Microm, Thermo Scientific, USA). The slices were adsorbed on SuperFrost Plus (Menzel, Germany) and stored at −80 °C until further use (30, 58). For [^18^F]PSS232 autoradiography, slices were thawed on ice and preconditioned with ice-cold buffer (pH 7.4) containing 30 mM HEPES, 1.2 mM MgCl_2_, 110 mM NaCl, 2.5 mM CaCl_2_, 5 mM KCl and 0.1% BSA. The tissue slices were dried and then incubated with 1 mL of [^18^F]PSS232 (1 nM) for 40 min at room temperature in a humidified chamber. For blockade conditions, 2-methyl-6-(phenylethynyl)-pyridine (MPEP; 1 μM) was added to the solution containing the radioligand. The slices were washed with ice-cold washing buffer (pH 7.4) containing 30 mM HEPES, 1.2 mM MgCl_2_, 110 mM NaCl, 2.5 mM CaCl_2_, 5 mM KCl, and ice-cold distilled water. For [^11^C]WMS1405B autoradiography, slices were thawed on ice and preconditioned. The tissue slices were dried and then incubated with 1 mL of [^11^C]WMS1405B (5 nM) for 15 min at room temperature in a humidified chamber. For blockade conditions, Ifenprodil (10 μM) was added to the solution containing the radioligand. The slices were washed with ice-cold washing buffer (pH 7.4) containing ice-cold distilled water. After drying, the slices were exposed to a phosphorimager plate (Fuji, Switzerland) for 30 min, and the film was scanned in a BAS5000 reader (Fuji, Japan).

### Immunohistochemistry, immunofluorescence staining, microscopy and image analysis

The paraffin-embedded fixed postmortem human brain tissues (from 34 AD patients and 31 NC subjects) were cut into 3 µm sections using a Leica microtome (Leica Microsystems, Germany). Hematoxylin and eosin (H&E) staining was performed according to routine procedures for each patient to provide anatomical information and to determine whether there were abnormalities in the brain. Immunochemical staining using antibodies against ionized calcium binding adaptor molecule 1 (Iba1) and Aβ17-24 (4G8) was performed. Immunofluorescence staining using antibodies against mGluR5, Aβ1-16 (6E10), P2X7R and phospho-Tau (AT-8) was performed. Paraffin-embedded fixed human brain tissue sections were incubated with primary antibodies overnight at 4°C with mild shaking (29). The detailed data and quantification of the levels of P2X7Rs in hippocampal tissue from NC and AD patients were obtained from our recent study (43). Here, the correlation between P2X7R and mGluR5 was assessed to determine the link between gliosis and mGluR5.

For immunofluorescence analysis of 3×Tg and arcAβ mouse brains, coronal mouse brain sections (40 mm) were cut around bregma 0 to −2 mm. Then, the sections were permeabilized and blocked in 5% normal donkey or goat serum and 1% Triton-PBS for one hour at room temperature with mild shaking. Free-floating tissue sections or paraffin-embedded sections were incubated with primary antibodies against 6E10 and mGluR5 overnight at 4°C (**STable 1**) (29). The next day, the sections were washed with PBS two times for 20 minutes and incubated with a suitable secondary antibody for 2 hours at room temperature. The sections were incubated for 15 minutes in 4′,6-diamidino-2-phenylindole (DAPI), washed two times for 10 minutes with PBS, and mounted with VECTASHIELD Vibrance Antifade Mounting Media (Vector Laboratories, Z J0215). The brain sections were imaged at ×20 magnification using an Axio Oberver Z1 slide scanner (Zeiss, Germany) using the same acquisition settings for all brain slices and at ×10 and ×63 magnification using a Leica SP8 confocal microscope (Leica, Germany). The images were analysed by a person blinded to the genotype using Qupath and ImageJ (NIH, U.S.A.).

For the human hippocampus, manual delineation of the adult human brain based on the Allen atlas (18) was performed using Qupath and ImageJ (NIH, U.S.A.). Regions of interest (ROIs) for CA1, CA2/3, the dentate gyrus (DG), the subiculum (SUB), and the entorhinal cortex (EC) were delineated. The mean fluorescence intensity was calculated after calibration for background intensity.

### RNA data analysis

The RNA sequencing data from the AD and NC groups used in this study were obtained from the scREAD database (https://bmbls.bmi.osumc.edu/scread/) (28), which comprises comprehensive datasets from GEO (Gene Expression Omnibus) (17) and Synapse, focusing specifically on human and mouse models of Alzheimer’s disease (AD). The specific datasets used in this study, along with their IDs, are as follows: Synapse IDs: syn18485175 and syn21125841; GEO IDs: GSE138852, GSE147528, GSE98969, GSE140510, GSE140399, GSE147495, GSE141044, GSE130626, GSE103334, GSE150358, GSE142853, GSE129308, and GSE146639. Data preprocessing involved removing duplicate entries, handling missing values, and standardizing gene expression values, particularly the log fold change (logFC) values, to ensure consistency across samples. The Mann-Whitney U test was used to compare gene expression levels between the AD and C groups across different cell types and brain regions (nonnormal data distributions). For comparisons between different cellular sources of GRM5 in the mouse brain and in the human brain, the Brain RNAseq Database was used (https://brainrnaseq.org/). The human data was based on (83), and the mouse data was based on (81).

### Proteomic profiling and pathway analysis

To further assess whether there is change in animal model with both amyloid and tau pathology, we conducted high-throughput quantitative proteomic analysis of hippocampal samples obtained from three 16-month-old 3×Tg mice and three age-matched wild-type (WT) mice. The analysis employed tandem mass tag (TMT) labelling, as previously described (34). Hippocampal tissues were collected from both 3×Tg mice and WT mice. Tissues were homogenized on ice for 10 minutes using lysis buffer (34) (**STable 2**). The resulting supernatant was obtained by centrifugation at 10,000×g for 30 minutes at 4°C. The supernatant was incubated with 100 mM tetraethylammonium bromide (TEAB) and 10 mM dithiothreitol (DTT) for 60 minutes at 55 °C. Subsequently, 35 mM iodoacetamide was added and incubated in the dark for 60 minutes. Acetonitrile (5 times volume, v/v) was added at −20°C for 3 hours. The sample was then centrifuged at 20,000×g for 30 minutes at 4 °C. The remaining sample underwent two additional incubations with 1 mL of 50% acetonitrile at −20 °C for 3 hours. To the protein precipitate, 0.1 mL of 100 mM TEAB was added, and the solution was mixed. The peptide/protein concentration was determined using the bicinchoninic acid (BCA) protein assay. The protein precipitate was combined with 100 mL of 100 mM TEAB, 1 mg/mL trypsin, and 1.0 mg of enzyme per 100 mg of protein. The mixture was incubated at 37°C for 4 hours, followed by the addition of additional trypsin and incubation at 37°C for 12 hours. The resulting peptide solution was centrifuged at 5,000 × g. The dried peptide powder was obtained using a freeze dryer. The enzymatic hydrolysis efficiency was assessed using LC MS. The peptide/protein concentration in the supernatant was determined by a BCA protein assay (34). For TMT labelling and LC MS analysis, the peptides were labelled using a TMT 6-plex kit following the manufacturer’s instructions. LC system analysis was subsequently performed. Extracted spectra from Proteome Discoverer (version 2.4.0.305) were searched with Sequest HT. Quantitative analysis of the spectra was carried out using Proteome Discoverer. The false positive rate (FDR) was set at ≤ 1% for both the protein and peptide levels (p value < 0.05; fold change > 1.2).

In addition, a comprehensive bioinformatics analysis was used to investigate the differentially expressed proteins and enriched signalling pathways as described earlier. We analysed the differentially expressed genes (DEPs) by using differential gene expression analysis and gene ontology (GO) analysis. Kyoto Encyclopedia of Genes and Genomes (KEGG) signalling pathway, kinase analysis, and KEGG signalling pathway analysis were performed using the *Mus musculus* (mouse) database (www.kegg.jp/kegg/pathway.html). GO analysis was performed using the GO database (http://www.geneontology.org/) with annotations based on the UniProt database (https://www.uniprot.org/taxonomy/10090). Protein-protein interaction (PPI) network analysis was performed for the DEPs using the STRING database for *Mus musculus* (mouse) (v11, stringdb.org). A volcano plot was generated using https://huygens.science.uva.nl/VolcaNoseR/.

### Statistics

All the statistical analyses were performed using Python and GraphPad Prism. The Shapiro-Wilk test was used to assess the normality of the distribution of various parameters in both groups. For the statistical analysis of differences between two groups, the nonparametric Mann-Whitney U test was used. Linear regression models were used to assess the impact of age and group status (AD vs. NC) on mGluR5 expression across brain regions, with age and group status being predictors and mGluR5 expression being the response variable. Differences in expression between groups were evaluated using p values for group coefficients to identify AD-related disparities. Nonparametric Spearman’s rank correlation analysis was performed between Braak stage, CERAD score, tau level, and Iba1, P2X7R and mGluR5 levels in the brain. Principal component analysis (PCA) and canonical correlation analysis (CCA) were conducted using R statistical software. PCA was utilized to explore the variance structure of the data. The datasets included two categories: tau proteins and mGluR5. Prior to PCA, data standardization was performed using the scale function in R to ensure that all variables contributed equally to the analysis by subtracting the mean and dividing by the standard deviation. PCA was executed using the prcomp function, which computes the covariance matrix, eigenvalues, and eigenvectors and extracts the principal components. The loadings of the first principal component were computed for each dataset to assess their contribution to variance. CCA was used to explore the relationships between mGluR5 expression levels and tau protein levels in the AD and NC datasets. Preprocessing steps involved removing observations with missing values. CCA was then performed separately for the AD and NC datasets with the objective of identifying linear combinations of mGluR5 and tau variables that maximized their correlation across brain regions. Canonical correlation coefficients and canonical loadings were calculated to quantify the strength and direction of the relationships between the two sets of variables, with canonical loadings representing the weights assigned to each variable in the linear combination.

## Results

### Demographics and pathological description and not affected by age, sex or APOE e4 allele status

The ages of the individuals in the NC group (82.2) were comparable to those in the AD group (82.4) and followed a normal statistical distribution (Shapiro-Wilk test). A greater percentage of females was noted in the AD group (88%) than in the NC group (58%). The prevalence of apolipoprotein E (APOE) ε4 carriers was significantly greater in the AD group (58.8%) than in the NC group (12.9%). H&E staining revealed no abnormalities in the brain tissue slices from the AD patients or NC subjects (data not shown). We used 4G8 immunohistochemical staining to map the Aβ distribution and AT-8 immunofluorescence staining to map the pathological phospho-Tau distribution in hippocampal slices. Amyloid-beta plaques and tau pathology were also observed in the brains of a proportion of NCs.

### Comparable hippocampal mGluR5 expression and GRM5 RNA levels in AD patients and NC subjects

We chose the hippocampus for the analysis of mGluR5 levels in AD and NC brains because it is the region with the most abundant mGluR5 in the human brain (2). We first quantified mGluR5 expression in subfields of the hippocampus in 34 AD patients and 30 NC subjects by using immunofluorescence staining and slide scanner microscopy. The mGluR5 levels decreased in the following order: the entorhinal cortex (EC), subiculum (SUB), CA3, CA1, and dentate gyrus (DG) in the AD group (**Fig. 1a**). In the NC group, the mGluR5 levels showed a similar decreasing trend across brain regions, with the highest levels observed in the SUB region, followed by the EC, CA3, CA1, and DG regions (**Figs. 1, 2**). Given that not all the patients had parahippocampal tissue blocks, we did not include parahippocampal tissue blocks in the data analysis. A similar level of mGluR5 was detected in all the hippocampal subfields analyzed (CA1, CA3, DG, Sub, and EC) in the AD patients compared to the NC subjects (**Fig. 1a**).

**Fig. 1.**
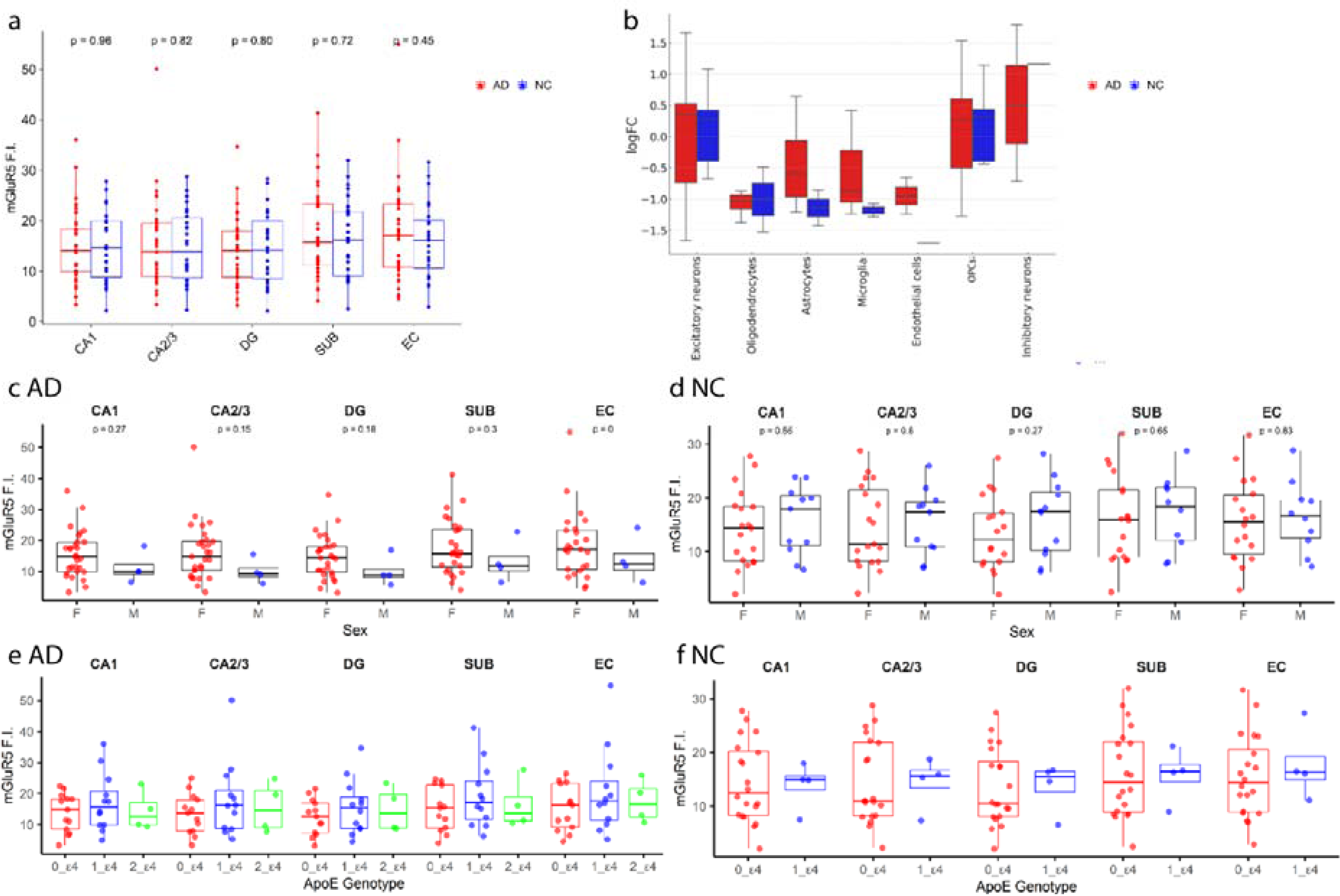
Distribution of mGluR5 in the hippocampus of AD patients and NC subjects. (a) No difference in mGluR5 levels between AD and NC groups. (b) No difference in GRM5 RNA expression in different cell types in the entorhinal cortex of AD and NC (n=25, n=13 for excitatory neurons; n=4, n=2 for oligodendrocytes; n=4, n=2 for astrocytes; n=3, n=2 for microglia; n=2, n=1 for endothelial cells; n=7, n=5 for oligodendrocyte precursor cells (OPCs); n=3, n=1 for inhibitory neurons). Data from the scREAD database (28) (https://bmbls.bmi.osumc.edu/scread/), which compiles comprehensive datasets from GEO (Gene Expression Omnibus) (17) and Synapse. logFC: log fold change. Boxplots showing the medians, quartiles, and outliers. (c) no difference in mGluR5 levels between male (n=4) and female AD patients (n=30) or (d) between male (n=18) and female NCs (n=12). (e) no difference in mGluR5 levels between APOE e4 carriers and noncarriers in the AD group or (f) in the NC group. DG, dentate gyrus; SUB, subiculum; EC, entorhinal cortex; CA, cornu ammonis.

**Fig. 2.**
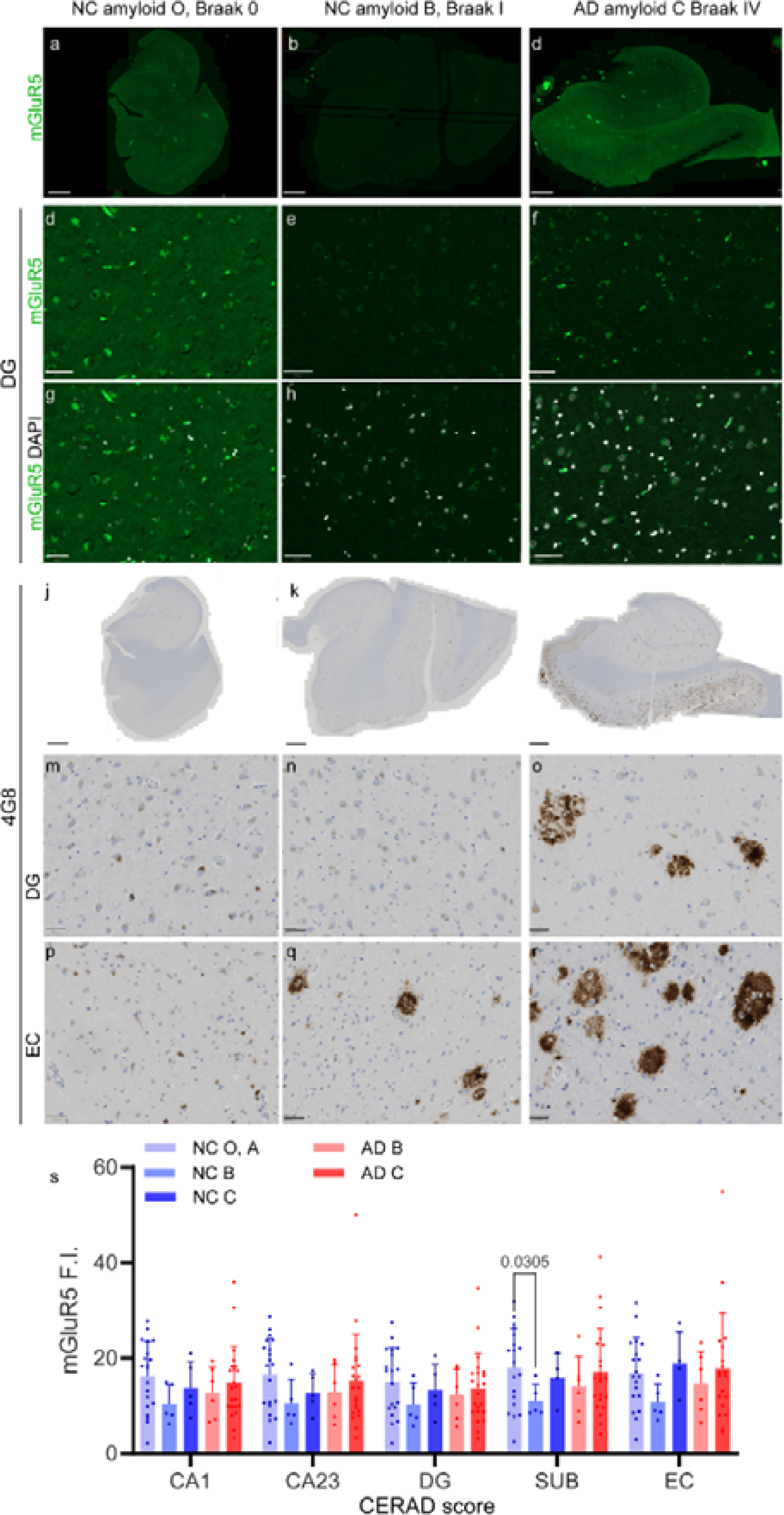
Reduced hippocampal mgluR5 in NC subjects with increased Aβ pathology. (**a-i**) Representative images of mGluR5 (green) staining and (**j-r**) 4G8 immunohistochemistry showing amyloid plaque deposits in the hippocampi of NC and AD patients with different amyloid levels (NCs #10-038, #04-015, and AD #01-116). Nuclei were counterstained with DAPI (white). (**s**) Quantification of mGluR5 fluorescence intensity in the CA1 region, CA2/3 region, DG, and EC. NC, Aβ score O-A, B, C (n=20, n=5, n=5). AD Aβ score B, C (n= 6, n= 23); Scale bar 2 mm (a-c, j-l), 50 mm (d-i, m-r). Nonparametric Mann-Whitney test.

Computation of RNA levels based on the existing dataset further revealed that there was no significant change in GRM5 levels in the entorhinal cortex between the AD group and the NC group; these changes included oligodendrocytes (U=4.0000, p=1.0000, AD n=4, NC n=2), astrocytes (U=6.0000, p=0.5333, AD n=4, NC n=2), microglia (U=5.0000, p=0.4000, AD n=3, NC n=2), endothelial cells (U=2.0000, p=0.6667, AD n=2, NC n=1), oligodendrocyte precursor cells (U=14.0000, p=0.6389, AD n=7, NC n=5), excitatory neurons (U=184.0000, p=0.5182, AD n=25, NC n=13), and inhibitory neurons (U=1.0000, p=1.0000, AD n=3, NC n=1) (**Fig. 1b**). Similar findings were found in the prefrontal cortex and in the superior frontal gyrus (BA8) (**SFig. 1**).

Next, we assessed the associations between hippocampal mGluR5 levels and age, APOE e4 status, and sex by using correlation and group analyses (**Fig. 1**). No correlation was detected between age and mGluR5 levels in the hippocampal subfields in the AD or NC groups (**Table 2**, linear transgression model). No difference in hippocampal mGluR5 levels was detected between females and males in either the AD (n=30 for females, n= 4 for males) or NC (n=18 for females, n=13 for males) groups (**Fig. 1c, d**). APOE ε4 is a genetic risk factor for sporadic AD. Here, we found no difference in hippocampal mGluR5 levels between APOE ε4 carriers and noncarriers in the NC and AD groups (**Figs. 1e, f**).

**Table 2.**
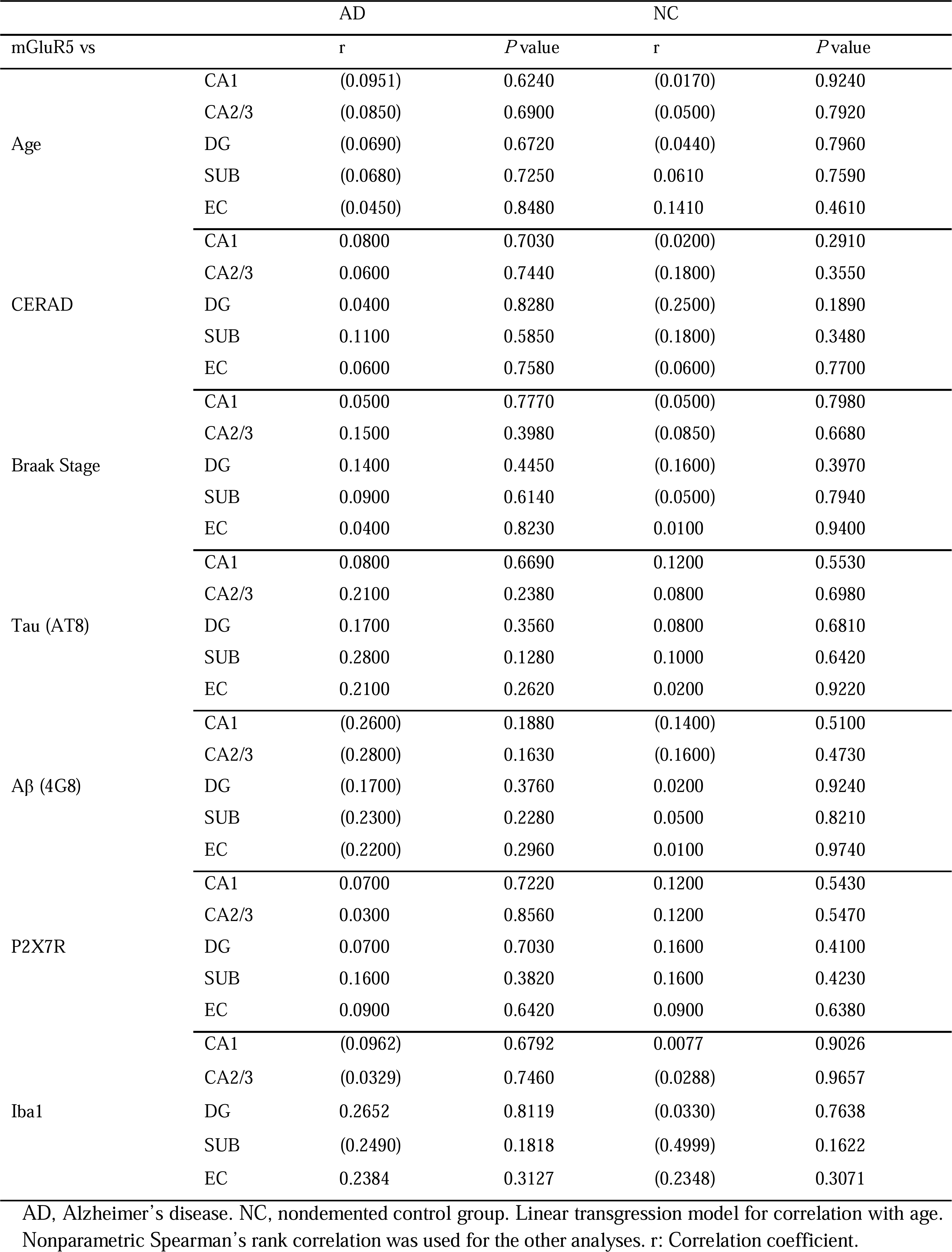
Correlation analysis in the Alzheimer’s disease and nondemented control groups.

### Reduced hippocampal mGluR5 levels with increased A**β** pathology in NCs

Next, we assessed the impact of Aβ accumulation on hippocampal mGluR5 levels in the NC and AD groups. An earlier study showed that Aβ oligomers, including mGluR5s, initiate synaptotoxicity following their interaction with the plasma membrane (73). Different types of amyloid plaques (CAAs, cored plaques, diffuse plaques) were observed, with a greater load in the hippocampus of AD patients than in that of NCs (**Fig. 2)**. Comparisons between AD patients with CERAD score B or C amyloid levels did not reveal significant differences. However, a lower mGluR5 level in the subiculum was detected in the NCs with an amyloid level of CERAD score B than in the NCs with scores of O and A (p = 0.0305, **Fig. 2 s**). Nonparametric Spearman’s rank analysis indicated no association between the amyloid level (both the CERAD score and % 4G8 positive area) and the mGluR5 level in the subfields of the hippocampus in the AD patients and NCs (**Table 2**). This finding indicated an association between amyloid pathology and the mGluR5 level in the hippocampus.

### Increased mGluR5 levels with increased tau in NCs and the association between mGluR5 and tau levels in the hippocampus

Next, we assessed the association between tau pathology and mGluR5 levels in the hippocampus. No significant difference was observed in the mGluR5 levels between AD patients with different tau levels (indicated by Braak stages II, III-IV, and V-VI). However, increased mGluR5 levels in the CA1 region, CA2/3 region, DG, and subiculum were detected in NC subjects in the Braak stage III-IV group compared with NC subjects in the Braak stage II group (CA1: p = 0.0076, CA2/3: p = 0.0372, DG: p = 0.0036, SUB: p = 0.0073) (**Fig. 3b,c, f, g, t**). This finding indicated a divergent influence of tau pathology on the mGluR5 level in the hippocampus, different from the effect of amyloid-beta pathology. Nonparametric Spearman’s rank analysis indicated no association between tau levels and mGluR5 levels in the subfields of the hippocampus in the AD patients and NCs (**Table 2**).

**Fig 3.**
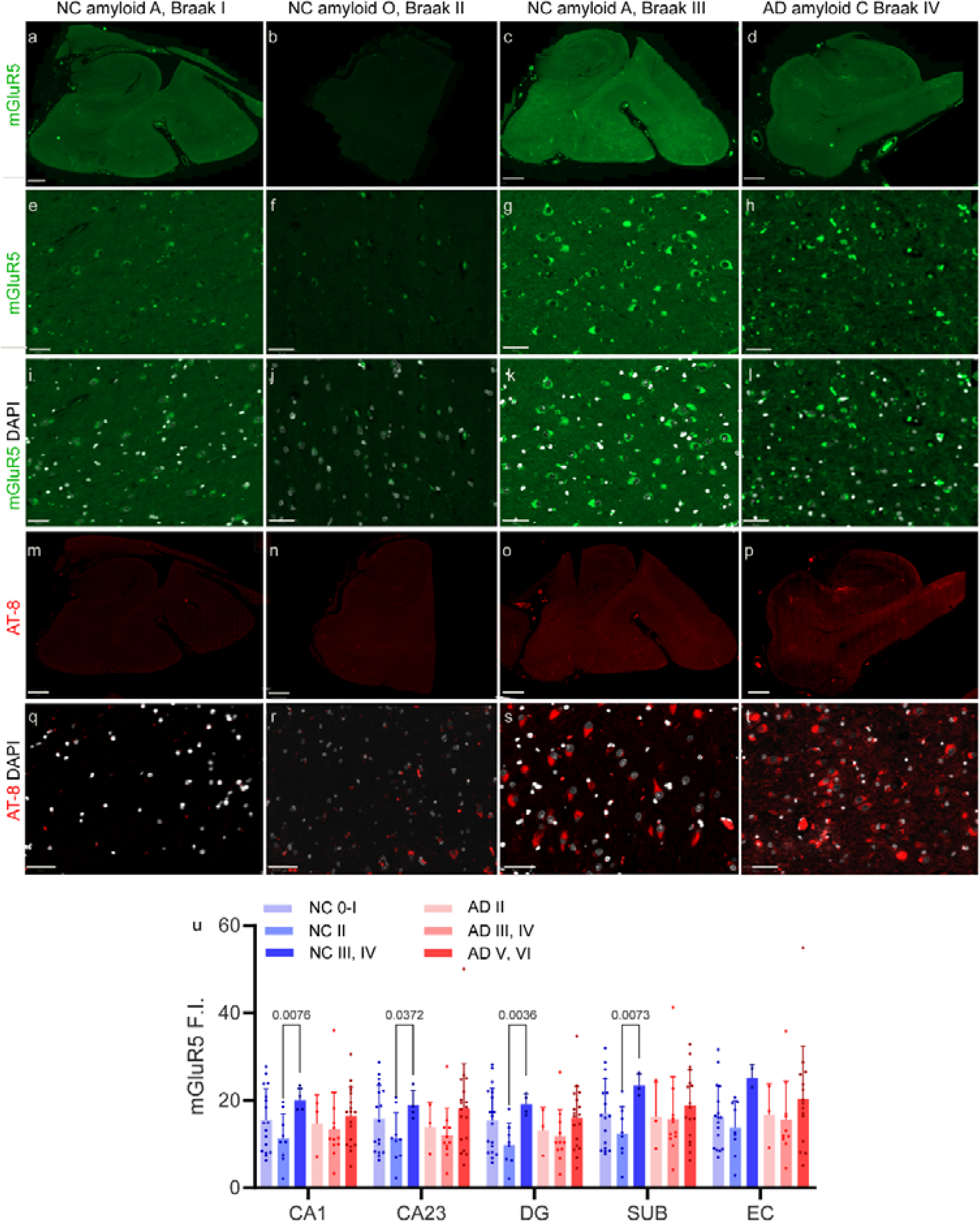
Increased mGluR5 in NC subjects with higher Braak tau stage. (**a-l**) Representative images of mGluR5 (green) and (**m-t**) phospho-Tau (AT8, red) staining in the hippocampi of NC (#08-103, #97-156, #05-073) and AD (#06-069). Increased hippocampal mgluR5 was associated with increased Braak stage (III-IV compared to II) in the NC brain. Zoom in in the entorhinal cortex (EC). Nuclei were counterstained with DAPI (white). (**u**) Quantification of mGluR5 fluorescence intensity in the CA1 region, CA2/3 region, dentate gyrus (DG), subiculum (SUB), and entorhinal cortex (EC) in the NC and AD groups. NC, Braak 0-I, II, III-IV (n=17, n= 9, n= 4); AD Braak II, III-IV, V-VI (n= 3, n=11, n=19). Scale bars: 2 mm (a-d, m-p) and 50 μm (e-l, q-t). Nonparametric Mann-Whitney test.

We further performed PCA and CCA to assess the distinct effects of tau on mGluR5 levels and associations between tau and mGluR5 levels in the hippocampus (**STable. 3, 4**). PCA of the AD and NC datasets. In the AD dataset, tau proteins exhibited positive loadings on the first principal component, with tau_SUB displaying the highest loading (0.337), indicating its predominant role in explaining variance within the AD group. Conversely, mGluR5 receptors demonstrated lower and more positive loading, suggesting that they contribute less to overall variance than do tau proteins. In contrast, in the NC dataset, tau proteins maintained positive loading, while mGluR5 receptors displayed negative loading, indicating an inverse relationship with tau protein expression patterns in the control group (**STable. 3**). CCA analysis revealed significant associations between mGluR5 expression in the EC region and tau protein levels in the CA2/3 region in the AD dataset. However, the mGluR5 expression level in the DG region was negatively associated with the tau protein level in the CA2/3 region. In the NC dataset, although associations were less pronounced, mGluR5 expression levels in the DG and SUB regions were positively associated with tau protein levels in the CA1 region (**STable. 4**).

### No associations between mGluR5 levels and Iba1 or P2X7R indicated gliosis in the hippocampus

An earlier study showed that mGluR5 plays a role in modulating neuroinflammation. An earlier study showed that P2X7R (which is expressed on both astrocytes and microglia) and mGlu5R crosstalk influence proinflammatory conditions in microglia (16, 75). Here, we assessed the associations of mGluR5 with Iba1 and P2X7R, which are indicative of gliosis, in the hippocampi of AD patients and HCs. In the hippocampus, which exhibited high levels of Iba1 staining, which is indicative of activated microglia (denser processes, **Fig 4n**), the mGluR5 level did not increase. Nonparametric Spearman’s rank analysis indicated no association between P2X7R levels and mGluR5 levels in the subfields of the hippocampus in the AD patients and NCs (**Table 2**). We further assessed the cellular source of GRM5 in the brains of humans and mice based on existing RNA sequencing data (81, 82). We found that there is a species difference in which neurons account for the majority (72%) of GRM5 source in the human brain, where as 10% for microglia and 4% for astrocytes (**Fig 4p**, Data computed from (82)), This is different from the pattern in mouse brain, where astroglial GRM5 is the majority (60%), followed by oliogodendroczte progenitor cell (21%), and neuron (15%) (**Fig 4q**, Data computed from (81)).

**Fig 4.**
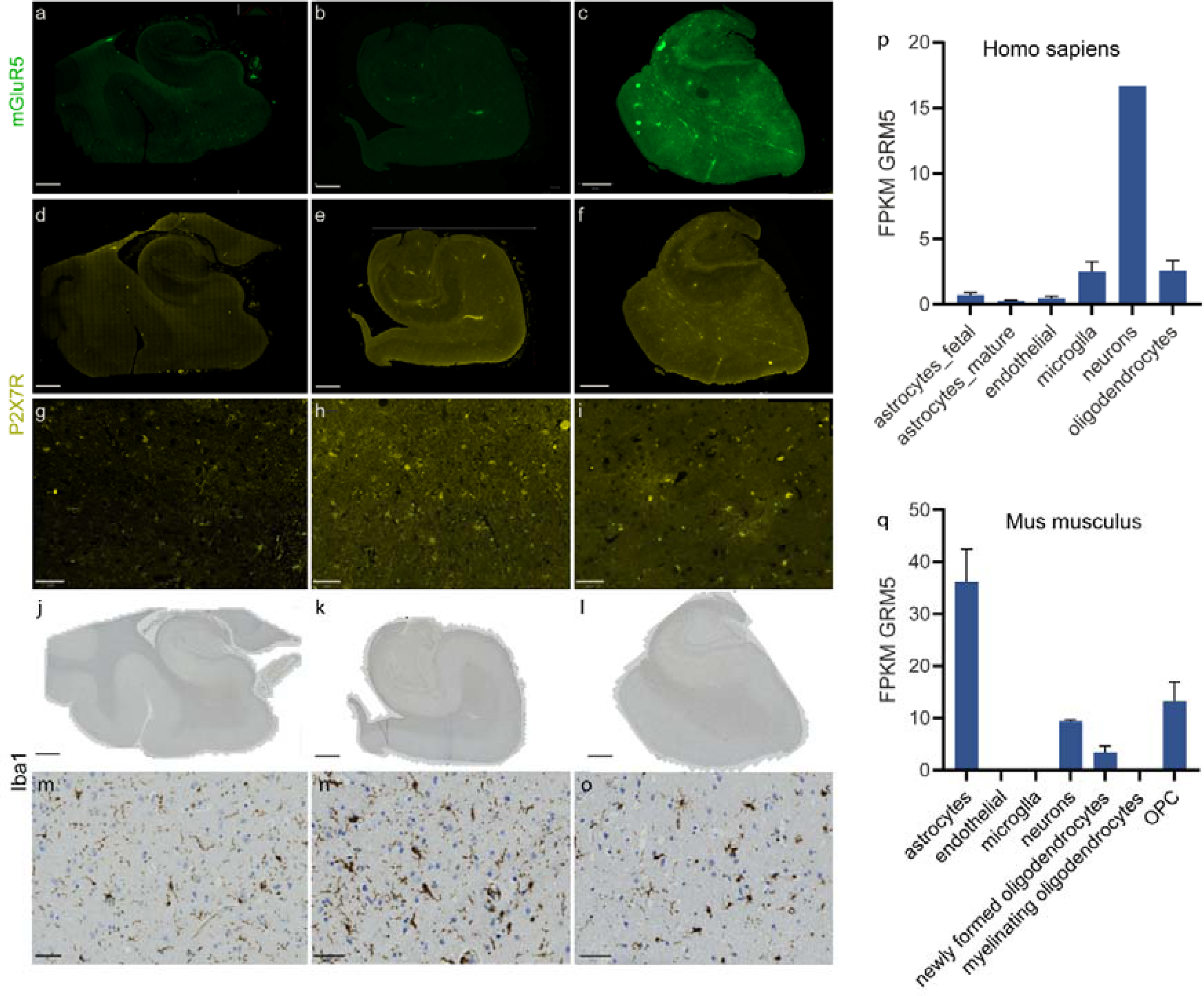
Distribution of microgliosis and mGluR5 in the hippocampus of NC subjects and AD patients. (**a-o**) Representative images of mGluR5 (green), corresponding P2X7R (yellow) and Iba1 (brown) staining in the NC (#09-022) and AD (#99-137, #99-098) hippocampi. Zoom in in the entorhinal cortex (EC). Scale bars: 2 mm (a-f, j-l) and 50 microns (g-i, m-o). (p, q) Different cellular sources of GRM5 RNA in the human brain and in the mouse brain. For comparisons between different cellular sources of GRM5 in the mouse brain and in the human brain, the Brain RNAseq Database was used (https://brainrnaseq.org/). Human data was based on (83), and mouse data was based on (81). FPKM: fragments per kilobase of transcript per million mapped reads.

### Comparable [^18^F]PSS232 binding in the brains of arcA**β** mice and nontransgenic littermates

We first assessed the changes in mGluR5 and GluN2B levels in amyloidosis mice that exhibited tau pathology. mGluR5 has been shown to be closely related to GluN2B in inducing downstream alterations (20). We examined GluN2B and mGluR5 levels by autoradiography using [^11^C]WMS1405B and [^18^F]PSS232, respectively, in nontransgenic littermate mice and arcAβ mice at 16 months of age. The nonspecific binding of both ligands is low in mouse brain tissue slices. No difference in [^18^F]PSS232 binding or [^11^C]WMS1405B binding was detected in the hippocampus or striatum between the arcAβ mice and their nontransgenic littermates (**Figs. 5a-c, SFig. 2**). We further performed immunofluorescence staining to assess whether there were differences in the levels of mGluR5 in the brain tissue of arcAβ mice and their nontransgenic littermates and whether mGluR5 expression and Aβ deposits were spatially associated. Abundant Aβ deposits were observed in the arcAβ mouse brain across the cortex, hippocampus and subcortical regions. We found that there was no clear difference in the immunofluorescence reactivity of mGluR5 between the brains of arcAβ mice and those of NTL mice (**Figs. 5d, e**). The mGluR5 level did not change depending on whether it was near or remote from the plaque in the hippocampus of the arcAβ mice (**Figs. 5e, f**).

**Fig. 5.**
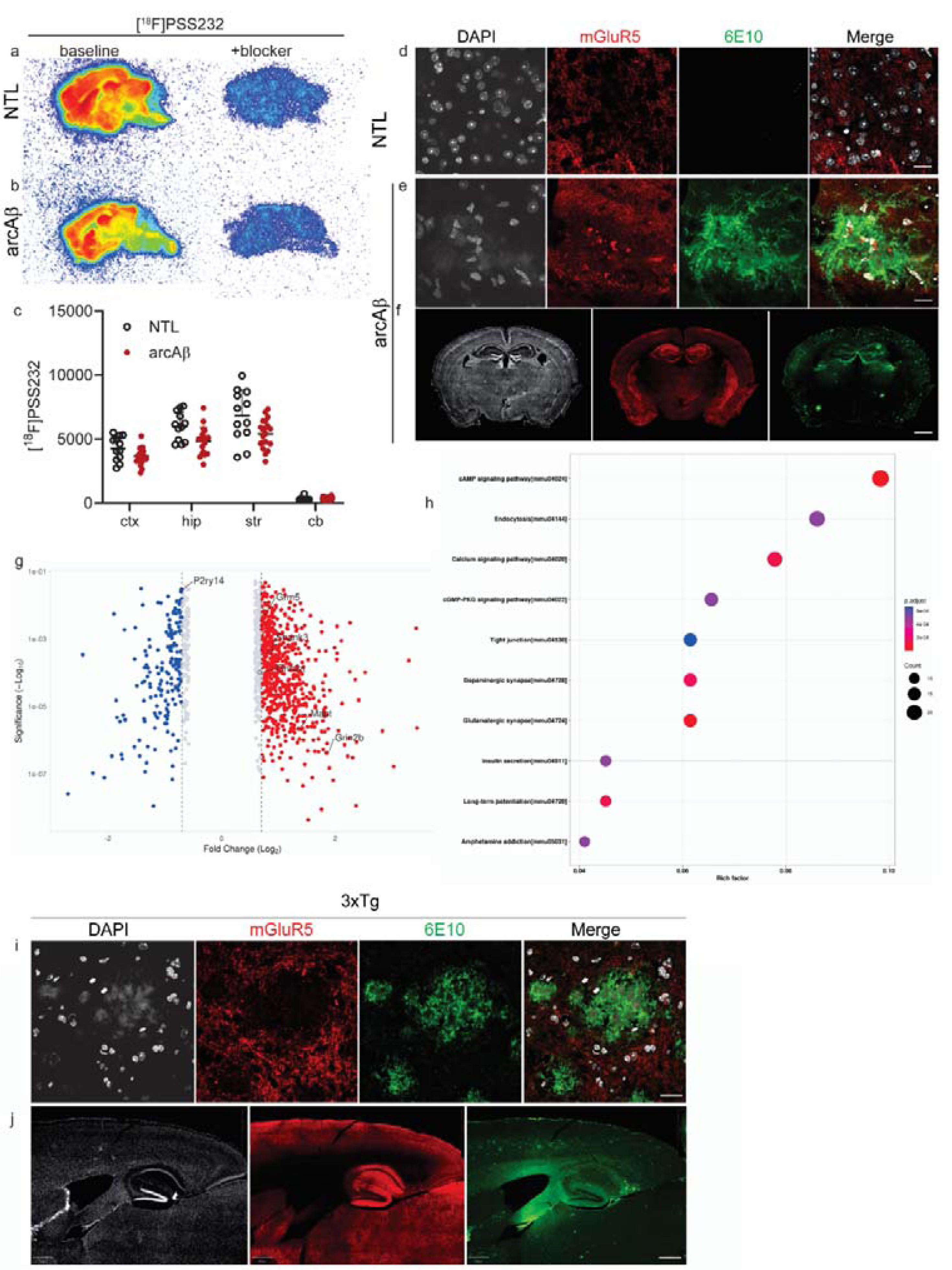
Analysis of mGluR5 expression and GRM5 levels in the brains of arcAβ mice and 3×Tg mice. (**a, b**) Representative [^18^F]PSS232 for mGluR5 autoradiographic images of sagittal brain sections of nontransgenic littermates (NTL) and arcAβ mice (baseline and in the presence of blocker). (**c**) Quantification of regional [^18^F]PSS232 binding in the cortex (ctx), hippocampus (hip), striatum (str), and cerebellum (cb). Nonparametric Mann-Whitney test, arcAβ vs. NTL. (**d-f**) Representative images of coronal brain slices from 16-month-old NTL mice and 16-month-old arcAβ mice stained for mGluR5 (red) and Aβ (6E10, green). (**g-h**) Kyoto Encyclopedia of Genes and Genomes (KEGG) analysis of the pathways associated with the differentially expressed proteins (DEPs) in the hippocampi of 16-month-old 3×Tg and NTL mice. (g) Volcano plots showing the log2-fold change (x-axis) and −log10 p value (y-axis) for all quantified proteins. (h) Enrichment analysis showing the top 10 enriched pathway terms. (**i, j**) Representative images of sagittal brain slices from 16-month-old 3×Tg mice stained for mGluR5 (red) and Aβ (6E10, green); nuclei were counterstained with DAPI (white). Scale bars: 20 μm (zoom in) and 800 μm.

### Increased GRM5 levels and glutamatergic pathway in the hippocampus of 3×Tg mice

To further understand the GRM5 alterations in AD, we selected 3×Tg mice with both Aβ and tau pathologies in the brain. Despite genetic drift, which showed a slower Aβ and tau accumulation in the brain compared to the original paper (27, 49), 3×Tg mice developed abundant Aβ and tau pathologies at 16 months of age according to a recent systematic characterization study as well as our own staining (**Fig. 6i, j**). GO and KEGG pathway enrichment analyses revealed increased levels of Shank3, Grm5 and MAPT, as well as upregulated glutamatergic pathways (top 7^th^) in hippocampal tissue from aged 3×Tg mice compared to that in hippocampal tissue from wild-type mice (**Fig. 5g, h, SFig 3, 4**). In addition, similar to that in the arcAβ mouse brain, the expression of mGluR5 in the hippocampus was greater than that in other brain regions in the 3×Tg mouse brain (**Fig. 5e, f, i, j**).

## Discussion

We provided evidence based on postmortem brain tissue from patients and animal models to demonstrate comparable mGluR5 and GRM5 levels in the hippocampus of AD patients. We revealed divergent and stage-dependent associations between mGluR5 and Aβ and between mGluR5 and tau pathologies in the hippocampus within the NC group. Our results are in line with earlier transcriptomic analysis of the prefrontal cortex of AD patients, which revealed altered expression of synaptic genes, including GRIN2A, GRIA1, GRIA2, GABRA1, GABRB2, GRM3 and SHANK2, (79), while GRM5 remained stable. The results from earlier *in vivo* PET studies revealed reduced [^11^C]ABP688 uptake in the hippocampus and amygdala in 9 AD patients and 10 NC subjects (70) and in 5 patients with behavioral variant frontotemporal dementia and 10 NC subjects (40). Reduced [^18^F]FPEB in the hippocampus in 16 mild cognitive impairment (MCI)/AD patients and 15 NC subjects (44) and [^18^F]PSS232 (19 AD patients and 16 NC subjects) (76). A recent [^18^F]PSS232 study (10 AD patients and 10 NC subjects) further showed that a reduction in mGluR5 availability in the hippocampus of AD was associated with PET imaging of Aβ deposits and glucose metabolism, and that this pattern was opposite to in the NC group (76). This negative correlation between Aβ and mGluR5 measured by [^18^F]PSS232 in the NC group (76) is in line with our results of lower hippocampal mGluR5 level in NCs with CERAD score II compared to NC with CERAD score of O and A. The relationship between tau and mGluR5 has not been reported *in vivo* by PET. Here, we found increased mGluR5 levels in NC subjects with Braak stage III-IV compared to NC subjects with Braak stage II. These findings indicate a complex, stage-dependent relationship between amyloid, tau and mGluR5.

There is a potential difference between the PET and *ex vivo* analyses of mGluR5 levels in the brain. One possible reason is that the PET tracers are antagonistic and agonist-based and might be more strongly associated with receptor activity than with *ex vivo* readouts. In addition, age-related tissue loss was implicated in one [^18^F]FPEB PET study: older age was associated with lower [^18^F]FPEB uptake (p= 0.026), whereas this association was not significant after gray matter masking or partial volume correction to account for age-related tissue loss (45).

Previous studies have shown that glutamate-dependent neuroglial calcium signaling differs between young and adult brains (66). Increased levels of mGluR5 levels have been reported in the hippocampal astrocytes of AD patients in proximity to Aβ plaques (15), colocalized with the nuclear accumulation of the p65 NF-kB subunit and increased CaNAα staining (41). However, according to the RNA sequencing data (**Fig. 4**), the expression of mGluR5 in the human brain is mainly neuronal (82). In addition, the fold changes in GRM5 levels were comparable in different cell types (neurons, astrocytes, microglia, etc.) in the entorhinal cortex between AD and NC groups. Therefore, we did not further investigate the neuronal/glial location of mGluR5 expression in the brains of patients with AD and HCs by using immunofluorescence staining. A link between mGluR5 and neuroinflammation has been reported: delayed mGluR5 activation has been shown to limit neuroinflammation (12) through the inhibition of NADPH oxidase (42), and neurodegeneration after traumatic brain injury (11). mGluR5 is also involved in AKT/PI3K signaling and the NF-κB pathway in methamphetamine-mediated increases in IL-6 and IL-8 expression in astrocytes (61). Agonist-dependent activation of mGluR5 inhibited alpha-synuclein-induced microglial inflammation and protected against neurotoxicity in the brain in a PD animal model (84). Using autoradiography, an earlier study revealed increased [^18^F]PSS232 binding in a lipopolysaccharide-induced neuroinflammation mouse model and in the brains of AD and amyotrophic lateral sclerosis patients (46). However, no prior study has directly investigated the association between the level of mGluR5 and neuroinflammation (gliosis) in the AD brain postmortem or *in vivo*. Here, we found that the mGluR5 level did not correlate with the P2X7R or Iba1 expression level, which is indicative of gliosis in the hippocampus of AD patients and NC subjects. In addition, we did not observe significant differences in the neuronal or glial expression of GRM5 (Fig. 1b) in the entorhinal cortex tissue of AD and NC.

Moreover, we found a species difference in the cellular location of GRM5, with glial cells dominant in the mouse brain and neuronal cells dominant in the human brain. This is important for developing imaging or therapeutic agents targeting mGluR5. Overexpression of mGluR5 has been shown in reactive astrocytes surrounding Aβ plaques in brain sections from APP/PS1 mice (14). Aβ oligomers led to the diffusional trapping and clustering of mGluR5s and an increase in ATP release following the activation of astroglial mGluR5s by their agonist (63). In addition, an earlier study in rats showed that astroglial mGluR5 expression in the hippocampus decreased with increasing age, with a greater decrease in the “a” splice variant than in the “b” splice variant (14).

The changes in mGluR5 in AD mouse models seem to depend on strain and stage according to earlier studies. In this study, we used an amyloidosis arcAβ mouse model with only amyloid deposits to determine the contribution of Aβ alone to mGluR5 levels and compared the results with those of 3×Tg mice with both amyloid and tau pathologies. We found no difference in mGluR5 levels in the hippocampus or striatum of arcAβ mice by using [^18^F]PSS232 autoradiography. Increase in the GRM5 level and enrichment of the glutamatergic pathway were observed via proteomics analysis of the hippocampal tissue of 3×Tg mice with both amyloid and tau pathologies. The network included APP and Prnp as well as genes linked to synaptic dysfunction in AD models, namely, Grm5, Fyn, and Ptk2B, as been also reported in previous studies (25). Increase in the mGluR5 surface expression (75%) but not mGluR5 total expression in the brain by using western blot was reported in 3×Tg mice at 9-months of age compared to control mice (4). A 12-week inhibition of mGluR5 using antagonist in 9-month-old 3×Tg mice shown improvement in the (4). Complex findings have been reported in other lines of AD mouse models: An earlier study showed increased mGluR5 expression on the cell surface but unaltered total mGluR5 levels in brain lysates of APP/PS1 mice at 9 months and 12 months, by using western blot (4, 22). Increased [^18^F]FPEB uptake was observed in 10-month-old APP/PS1 mice and age-matched NTL mice (39), while another study showed slightly greater (not significant) [^18^F]FPEB uptake in the brain of 10-month-old APP/PS1 mice (74). mGluR5 contribution to AD neuropathology has shown a disease stage-dependent pattern: blockade of mGluR5 for 24 weeks but not for 36 weeks was shown to reduce Aβ pathology, neuroinflammation and the cognitive impairment in 6-month-old APP/PS1 mice (3), whereas the effect was abolished in 15 month old APP/PS1 mice. Reduced levels of mGluR5 have been reported in the cortex, hippocampus and striatum by using [^18^F]FPEB PET and histology in 5×FAD mice aged 9 months compared to 3 months (38). Furthermore, by immunoblotting and immunofluorescence staining, [^11^C]ABP688 PET revealed elevated mGluR5 in 16-month-old tg-ArcSwe mice compared with control mice (19). A reduction in mGluR5 levels has been reported by using [^11^C]ABP688 PET and western blot in a 6×Tg mouse model of AD (31). Earlier studies have shown increased mGluR5 levels in the hippocampus of α-syn/APP tg mice and that mGluR5 mediated the toxicity of increased calcium flux and the vulnerability of hippocampal neurons to alpha-synuclein and Aβ (50). [^11^C]ABP688 uptake in the brain of P301L tau mouse model (rTg4510) was found to be unaltered relative to that in nontransgenic mice at 2 months. However, the expression of Grm4 gradually decreases in parallel with progressive brain atrophy at 5-6 and 8-9 months in an rTg4510 mouse model of tauopathy (62).

There are several limitations in our study. First, there was a greater percentage of females in the AD group than in the NC group. Although Aβ oligomers have been shown to induce pathophysiological mGluR5 signaling in AD mice in a sex-selective manner (1). We found that sex did not influence the hippocampal mGluR5 expression levels in AD and NC groups. Second, the clinical history, Mini-Mental State Examination (MMSE) scores and smoking habits (6) of the AD patients and NC subjects were not known, which might be related to mGluR5 levels.

## Conclusion

In the present study, we found comparable hippocampal mGluR5 levels between AD patients and NC individuals. The hippocampal level of mGluR5 is differentially influenced by amyloid and tau but not by age, sex or APOE e4 genotype. According to earlier mechanistic studies, the clustering of mGluR5 rather than the amount of mGluR5 might be the major target for therapeutic intervention (57).

## Acknowledgments

The authors acknowledge the NBB for providing postmortem human brain tissue; the authors acknowledge the Center for Microscopy and Image Analysis (ZMB), Marine Bruttin and Daniel Schuppli at the Institute for Regenerative Medicine, University of Zurich, and Ms. Claudia Keller at the Institute for Pharmaceutical Sciences for their technical assistance.

## Author contributions

RN conceived and designed the study. JW performed the statistical programming and data analysis. SS performed the quantification of microscopy data. YK performed the proteomics and data analysis. CM performed the confocal microscopy. RN performed the autoradiography. LM performed the radiolabeling and quality control of the radioligands. JW and RN wrote the first draft. All the authors contributed to the revision of the manuscript. All the authors have read and approved the final manuscript.

## References

1. Abd-Elrahman KS, Albaker A, de Souza JM, Ribeiro FM, Schlossmacher MG, Tiberi M, Hamilton A, Ferguson SSG (2020) Aβ oligomers induce pathophysiological mGluR5 signaling in Alzheimer’s disease model mice in a sex-selective manner. Sci Signal.13(662).

2. Abd-Elrahman KS, Ferguson SSG (2022) Noncanonical Metabotropic Glutamate Receptor 5 Signaling in Alzheimer’s Disease. Annu Rev Pharmacol Toxicol.62:235–54.

3. Abd-Elrahman KS, Hamilton A, Albaker A, Ferguson SSG (2020) mGluR5 Contribution to Neuropathology in Alzheimer Mice Is Disease Stage-Dependent. ACS Pharmacol Transl Sci.3(2):334–44.

4. Abd-Elrahman KS, Hamilton A, Vasefi M, Ferguson SSG (2018) Autophagy is increased following either pharmacological or genetic silencing of mGluR5 signaling in Alzheimer’s disease mouse models. Mol Brain.11(1):19.

5. Ahmed H, Wallimann R, Haider A, Hosseini V, Gruber S, Robledo M, Nguyen TAN, Herde AM, Iten I, Keller C, Vogel V, Schibli R, Wünsch B, Mu L, Ametamey SM (2021) Preclinical Development of (18)F-OF-NB1 for Imaging GluN2B-Containing N-Methyl-d-Aspartate Receptors and Its Utility as a Biomarker for Amyotrophic Lateral Sclerosis. J Nucl Med.62(2):259–65.

6. Akkus F, Ametamey SM, Treyer V, Burger C, Johayem A, Umbricht D, Gomez Mancilla B, Sovago J, Buck A, Hasler G (2013) Marked global reduction in mGluR5 receptor binding in smokers and ex-smokers determined by [11C]ABP688 positron emission tomography. Proc Natl Acad Sci U S A.110(2):737–42.

7. Albasanz JL, Dalfó E, Ferrer I, Martín M (2005) Impaired metabotropic glutamate receptor/phospholipase C signaling pathway in the cerebral cortex in Alzheimer’s disease and dementia with Lewy bodies correlates with stage of Alzheimer’s-disease-related changes. Neurobiol Dis.20(3):685–93.

8. Ametamey SM, Kessler LJ, Honer M, Wyss MT, Buck A, Hintermann S, Auberson YP, Gasparini F, Schubiger PA (2006) Radiosynthesis and preclinical evaluation of 11C-ABP688 as a probe for imaging the metabotropic glutamate receptor subtype 5. J Nucl Med.47(4):698–705.

9. Braak H, Braak E (1991) Neuropathological stageing of Alzheimer-related changes. Acta Neuropathol.82(4):239–59.

10. Bruno V, Battaglia G, Copani A, D’Onofrio M, Di Iorio P, De Blasi A, Melchiorri D, Flor PJ, Nicoletti F (2001) Metabotropic glutamate receptor subtypes as targets for neuroprotective drugs. J Cereb Blood Flow Metab.21(9):1013–33.

11. Byrnes KR, Loane DJ, Stoica BA, Zhang J, Faden AI (2012) Delayed mGluR5 activation limits neuroinflammation and neurodegeneration after traumatic brain injury. J Neuroinflammation.9:43.

12. Byrnes KR, Stoica B, Loane DJ, Riccio A, Davis MI, Faden AI (2009) Metabotropic glutamate receptor 5 activation inhibits microglial associated inflammation and neurotoxicity. Glia.57(5):550–60.

13. Cai L, Liow JS, Morse CL, Telu S, Davies R, Frankland MP, Zoghbi SS, Cheng K, Hall MD, Innis RB, Pike VW (2020) Evaluation of (11)C-NR2B-SMe and Its Enantiomers as PET Radioligands for Imaging the NR2B Subunit Within the NMDA Receptor Complex in Rats. J Nucl Med.61(8):1212–20.

14. Cai Z, Schools GP, Kimelberg HK (2000) Metabotropic glutamate receptors in acutely isolated hippocampal astrocytes: developmental changes of mGluR5 mRNA and functional expression. Glia.29(1):70–80.

15. Casley CS, Lakics V, Lee H-g, Broad LM, Day TA, Cluett T, Smith MA, O’Neill MJ, Kingston AE (2009) Up-regulation of astrocyte metabotropic glutamate receptor 5 by amyloid-β peptide. Brain Research.1260:65–75.

16. Chisari M, Barraco M, Bucolo C, Ciranna L, Sortino MA (2023) Purinergic ionotropic P2X7 and metabotropic glutamate mGlu(5) receptors crosstalk influences pro-inflammatory conditions in microglia. Eur J Pharmacol.938:175389.

17. Clough E, Barrett T (2016) The Gene Expression Omnibus Database. Methods Mol Biol.1418:93–110.

18. Ding SL, Royall JJ, Sunkin SM, Ng L, Facer BA, Lesnar P, Guillozet-Bongaarts A, McMurray B, Szafer A, Dolbeare TA, Stevens A, Tirrell L, Benner T, Caldejon S, Dalley RA, Dee N, Lau C, Nyhus J, Reding M, Riley ZL, Sandman D, Shen E, van der Kouwe A, Varjabedian A, Wright M, Zöllei L, Dang C, Knowles JA, Koch C, Phillips JW, Sestan N, Wohnoutka P, Zielke HR, Hohmann JG, Jones AR, Bernard A, Hawrylycz MJ, Hof PR, Fischl B, Lein ES (2016) Comprehensive cellular-resolution atlas of the adult human brain. J Comp Neurol.524(16):3127–481.

19. Fang XT, Eriksson J, Antoni G, Yngve U, Cato L, Lannfelt L, Sehlin D, Syvänen S (2017) Brain mGluR5 in mice with amyloid beta pathology studied with in vivo [(11)C]ABP688 PET imaging and ex vivo immunoblotting. Neuropharmacology.113(Pt A):293–300.

20. Gu L, Luo WY, Xia N, Zhang JN, Fan JK, Yang HM, Wang MC, Zhang H (2022) Upregulated mGluR5 induces ER stress and DNA damage by regulating the NMDA receptor subunit NR2B. J Biochem.171(3):349–59.

21. Haas LT, Salazar SV, Smith LM, Zhao HR, Cox TO, Herber CS, Degnan AP, Balakrishnan A, Macor JE, Albright CF, Strittmatter SM (2017) Silent Allosteric Modulation of mGluR5 Maintains Glutamate Signaling while Rescuing Alzheimer’s Mouse Phenotypes. Cell Rep.20(1):76–88.

22. Hamilton A, Esseltine JL, DeVries RA, Cregan SP, Ferguson SS (2014) Metabotropic glutamate receptor 5 knockout reduces cognitive impairment and pathogenesis in a mouse model of Alzheimer’s disease. Mol Brain.7:40.

23. Hamilton A, Vasefi M, Vander Tuin C, McQuaid RJ, Anisman H, Ferguson SS (2016) Chronic Pharmacological mGluR5 Inhibition Prevents Cognitive Impairment and Reduces Pathogenesis in an Alzheimer Disease Mouse Model. Cell Rep.15(9):1859–65.

24. Hannan AJ, Blakemore C, Katsnelson A, Vitalis T, Huber KM, Bear M, Roder J, Kim D, Shin HS, Kind PC (2001) PLC-beta1, activated via mGluRs, mediates activity-dependent differentiation in cerebral cortex. Nat Neurosci.4(3):282–8.

25. Heiss JK, Barrett J, Yu Z, Haas LT, Kostylev MA, Strittmatter SM (2017) Early Activation of Experience-Independent Dendritic Spine Turnover in a Mouse Model of Alzheimer’s Disease. Cereb Cortex.27(7):3660–74.

26. Jakobsson JE, Gourni E, Khanapur S, Brito B, Riss PJ (2019) Synthesis and Characterization in Rodent Brain of the Subtype-Selective NR2B NMDA Receptor Ligand [(11)C]Ro04-5595 as a Potential Radiotracer for Positron Emission Tomography. ACS Omega.4(6):9925–31.

27. Javonillo DI, Tran KM, Phan J, Hingco E, Kramár EA, da Cunha C, Forner S, Kawauchi S, Milinkeviciute G, Gomez-Arboledas A, Neumann J, Banh CE, Huynh M, Matheos DP, Rezaie N, Alcantara JA, Mortazavi A, Wood MA, Tenner AJ, MacGregor GR, Green KN, LaFerla FM (2021) Systematic Phenotyping and Characterization of the 3xTg-AD Mouse Model of Alzheimer’s Disease. Front Neurosci.15:785276.

28. Jiang J, Wang C, Qi R, Fu H, Ma Q (2020) scREAD: A Single-Cell RNA-Seq Database for Alzheimer’s Disease. iScience.23(11):101769.

29. Kecheliev V, Boss L, Maheshwari U, Konietzko U, Keller A, Razansky D, Nitsch RM, Klohs J, Ni R (2023) Aquaporin 4 is differentially increased and dislocated in association with tau and amyloid-beta. Life Sciences.121593.

30. Kecheliev V, Spinelli F, Herde A, Haider A, Mu L, Klohs J, Ametamey SM, Ni R (2022) Evaluation of cannabinoid type 2 receptor expression and pyridine-based radiotracers in brains from a mouse model of Alzheimer’s disease. Frontiers in Aging Neuroscience.14.

31. Kim Y, Kim J, Kang S, Chang KA (2023) Depressive-like Behaviors Induced by mGluR5 Reduction in 6xTg in Mouse Model of Alzheimer’s Disease. Int J Mol Sci.24(16).

32. Knopman DS, Amieva H, Petersen RC, Chételat G, Holtzman DM, Hyman BT, Nixon RA, Jones DT (2021) Alzheimer disease. Nat Rev Dis Primers.7(1):33.

33. Kong Y, Cao L, Wang J, Zhuang J, Liu Y, Bi L, Qiu Y, Hou Y, Huang Q, Xie F, Yang Y, Shi K, Rominger A, Guan Y, Jin H, Ni R (2024) Increased Cerebral Level of P2X7R in a Tauopathy Mouse Model by PET Using [(18)F]GSK1482160. ACS Chem Neurosci.

34. Kong Y, Cao L, Xie F, Wang X, Zuo C, Shi K, Rominger A, Huang Q, Xiao J, Jiang D, Guan Y, Ni R (2024) Reduced SV2A and GABAA receptor levels in the brains of type 2 diabetic rats revealed by [18F]SDM-8 and [18F]flumazenil PET. Biomedicine & Pharmacotherapy.172:116252.

35. Kong Y, Huang L, Li W, Liu X, Zhou Y, Liu C, Zhang S, Xie F, Zhang Z, Jiang D, Zhou W, Ni R, Zhang C, Sun B, Wang J, Guan Y (2021) The Synaptic Vesicle Protein 2A Interacts With Key Pathogenic Factors in Alzheimer’s Disease: Implications for Treatment. Frontiers in cell and developmental biology.9:609908-.

36. Kong Y, Maschio CA, Shi X, Xie F, Zuo C, Konietzko U, Shi K, Rominger A, Xiao J, Huang Q, Nitsch RM, Guan Y, Ni R (2024) Relationship Between Reactive Astrocytes, by [(18)F]SMBT-1 Imaging, with Amyloid-Beta, Tau, Glucose Metabolism, and TSPO in Mouse Models of Alzheimer’s Disease. Mol Neurobiol.

37. Krämer SD, Betzel T, Mu L, Haider A, Herde AM, Boninsegni AK, Keller C, Szermerski M, Schibli R, Wünsch B, Ametamey SM (2018) Evaluation of (11)C-Me-NB1 as a Potential PET Radioligand for Measuring GluN2B-Containing NMDA Receptors, Drug Occupancy, and Receptor Cross Talk. J Nucl Med.59(4):698–703.

38. Lee M, Lee HJ, Jeong YJ, Oh SJ, Kang KJ, Han SJ, Nam KR, Lee YJ, Lee KC, Ryu YH, Hyun IY, Choi JY (2019) Age dependency of mGluR5 availability in 5xFAD mice measured by PET. Neurobiol Aging.84:208–16.

39. Lee M, Lee HJ, Park IS, Park JA, Kwon YJ, Ryu YH, Kim CH, Kang JH, Hyun IY, Lee KC, Choi JY (2018) Aβ pathology downregulates brain mGluR5 density in a mouse model of Alzheimer. Neuropharmacology.133:512–7.

40. Leuzy A, Zimmer ER, Dubois J, Pruessner J, Cooperman C, Soucy JP, Kostikov A, Schirmaccher E, Désautels R, Gauthier S, Rosa-Neto P (2016) In vivo characterization of metabotropic glutamate receptor type 5 abnormalities in behavioral variant FTD. Brain Struct Funct.221(3):1387–402.

41. Lim D, Iyer A, Ronco V, Grolla AA, Canonico PL, Aronica E, Genazzani AA (2013) Amyloid beta deregulates astroglial mGluR5-mediated calcium signaling via calcineurin and Nf-kB. Glia.61(7):1134–45.

42. Loane DJ, Stoica BA, Pajoohesh-Ganji A, Byrnes KR, Faden AI (2009) Activation of metabotropic glutamate receptor 5 modulates microglial reactivity and neurotoxicity by inhibiting NADPH oxidase. J Biol Chem.284(23):15629–39.

43. Maschio C, Wang J, Maheshwari U, Keller A, Rominger A, Konietzko U, Hock C, Nitsch RM, Nordberg A, Ni R (2024) Hippocampal purinergic P2X7 receptor level is increased in Alzheimer′s disease patients, and associated with amyloid and tau pathologies. bioRxiv.

44. Mecca AP, McDonald JW, Michalak HR, Godek TA, Harris JE, Pugh EA, Kemp EC, Chen MK, Salardini A, Nabulsi NB, Lim K, Huang Y, Carson RE, Strittmatter SM, van Dyck CH (2020) PET imaging of mGluR5 in Alzheimer’s disease. Alzheimers Res Ther.12(1):15.

45. Mecca AP, Rogers K, Jacobs Z, McDonald JW, Michalak HR, DellaGioia N, Zhao W, Hillmer AT, Nabulsi N, Lim K, Ropchan J, Huang Y, Matuskey D, Esterlis I, Carson RE, van Dyck CH (2021) Effect of age on brain metabotropic glutamate receptor subtype 5 measured with [(18)F]FPEB PET. Neuroimage.238:118217.

46. Müller Herde A, Schibli R, Weber M, Ametamey SM (2019) Metabotropic glutamate receptor subtype 5 is altered in LPS-induced murine neuroinflammation model and in the brains of AD and ALS patients. Eur J Nucl Med Mol Imaging.46(2):407–20.

47. Ni R, Müller Herde A, Haider A, Keller C, Louloudis G, Vaas M, Schibli R, Ametamey SM, Klohs J, Mu L (2021) In vivo Imaging of Cannabinoid Type 2 Receptors: Functional and Structural Alterations in Mouse Model of Cerebral Ischemia by PET and MRI. Molecular Imaging and Biology.

48. Niswender CM, Conn PJ (2010) Metabotropic glutamate receptors: physiology, pharmacology, and disease. Annu Rev Pharmacol Toxicol.50:295–322.

49. Oddo S, Caccamo A, Shepherd JD, Murphy MP, Golde TE, Kayed R, Metherate R, Mattson MP, Akbari Y, LaFerla FM (2003) Triple-transgenic model of Alzheimer’s disease with plaques and tangles: intracellular Abeta and synaptic dysfunction. Neuron.39(3):409–21.

50. Overk CR, Cartier A, Shaked G, Rockenstein E, Ubhi K, Spencer B, Price DL, Patrick C, Desplats P, Masliah E (2014) Hippocampal neuronal cells that accumulate α-synuclein fragments are more vulnerable to Aβ oligomer toxicity via mGluR5--implications for dementia with Lewy bodies. Mol Neurodegener.9:18.

51. Paoletti P, Bellone C, Zhou Q (2013) NMDA receptor subunit diversity: impact on receptor properties, synaptic plasticity and disease. Nat Rev Neurosci.14(6):383–400.

52. Pascoal TA, Benedet AL, Ashton NJ, Kang MS, Therriault J, Chamoun M, Savard M, Lussier FZ, Tissot C, Karikari TK, Ottoy J, Mathotaarachchi S, Stevenson J, Massarweh G, Schöll M, de Leon MJ, Soucy JP, Edison P, Blennow K, Zetterberg H, Gauthier S, Rosa-Neto P (2021) Microglial activation and tau propagate jointly across Braak stages. Nat Med.

53. Pillai RL, Tipre DN (2016) Metabotropic glutamate receptor 5 - a promising target in drug development and neuroimaging. Eur J Nucl Med Mol Imaging.43(6):1151–70.

54. Pinky PD, Pfitzer JC, Senfeld J, Hong H, Bhattacharya S, Suppiramaniam V, Qureshi I, Reed MN (2023) Recent Insights on Glutamatergic Dysfunction in Alzheimer’s Disease and Therapeutic Implications. Neuroscientist.29(4):461–71.

55. Rammes G, Mattusch C, Wulff M, Seeser F, Kreuzer M, Zhu K, Deussing JM, Herms J, Parsons CG (2017) Involvement of GluN2B subunit containing N-methyl-d-aspartate (NMDA) receptors in mediating the acute and chronic synaptotoxic effects of oligomeric amyloid-beta (Aβ) in murine models of Alzheimer’s disease (AD). Neuropharmacology.123:100–15.

56. Ren W, Li L, Zhang J, Vaas M, Klohs J, Ripoll J, Wolf M, Ni R, Rudin M (2022) Non-invasive visualization of amyloid-beta deposits in Alzheimer amyloidosis mice using magnetic resonance imaging and fluorescence molecular tomography. Biomed Opt Express.13(7):3809–22.

57. Renner M, Lacor PN, Velasco PT, Xu J, Contractor A, Klein WL, Triller A (2010) Deleterious effects of amyloid beta oligomers acting as an extracellular scaffold for mGluR5. Neuron.66(5):739–54.

58. Rodriguez-Vieitez E, Ni R, Gulyás B, Tóth M, Häggkvist J, Halldin C, Voytenko L, Marutle A, Nordberg A (2015) Astrocytosis precedes amyloid plaque deposition in Alzheimer APPswe transgenic mouse brain: a correlative positron emission tomography and in vitro imaging study. Eur J Nucl Med Mol Imaging.42(7):1119–32.

59. Sathe G, Mangalaparthi KK, Jain A, Darrow J, Troncoso J, Albert M, Moghekar A, Pandey A (2020) Multiplexed Phosphoproteomic Study of Brain in Patients with Alzheimer’s Disease and Age-Matched Cognitively Healthy Controls. Omics.24(4):216–27.

60. Sephton SM, Herde AM, Mu L, Keller C, Rüdisühli S, Auberson Y, Schibli R, Krämer SD, Ametamey SM (2015) Preclinical evaluation and test-retest studies of [(18)F]PSS232, a novel radioligand for targeting metabotropic glutamate receptor 5 (mGlu5). Eur J Nucl Med Mol Imaging.42(1):128–37.

61. Shah A, Silverstein PS, Singh DP, Kumar A (2012) Involvement of metabotropic glutamate receptor 5, AKT/PI3K signaling and NF-κB pathway in methamphetamine-mediated increase in IL-6 and IL-8 expression in astrocytes. J Neuroinflammation.9:52.

62. Shimojo M, Takuwa H, Takado Y, Tokunaga M, Tsukamoto S, Minatohara K, Ono M, Seki C, Maeda J, Urushihata T, Minamihisamatsu T, Aoki I, Kawamura K, Zhang MR, Suhara T, Sahara N, Higuchi M (2020) Selective Disruption of Inhibitory Synapses Leading to Neuronal Hyperexcitability at an Early Stage of Tau Pathogenesis in a Mouse Model. J Neurosci.40(17):3491–501.

63. Shrivastava AN, Kowalewski JM, Renner M, Bousset L, Koulakoff A, Melki R, Giaume C, Triller A (2013) β-amyloid and ATP-induced diffusional trapping of astrocyte and neuronal metabotropic glutamate type-5 receptors. Glia.61(10):1673–86.

64. Soares C, Da Ros LU, Machado LS, Rocha A, Lazzarotto G, Carello-Collar G, De Bastiani MA, Ferrari-Souza JP, Lussier FZ, Souza DO, Rosa-Neto P, Pascoal TA, Bellaver B, Zimmer ER (2024) The glutamatergic system in Alzheimer’s disease: a systematic review with meta-analysis. In: Mol Psychiatry, © 2024. The Author(s), under exclusive licence to Springer Nature Limited.: England.

65. Spurrier J, Nicholson L, Fang XT, Stoner AJ, Toyonaga T, Holden D, Siegert TR, Laird W, Allnutt MA, Chiasseu M, Brody AH, Takahashi H, Nies SH, Pérez-Cañamás A, Sadasivam P, Lee S, Li S, Zhang L, Huang YH, Carson RE, Cai Z, Strittmatter SM (2022) Reversal of synapse loss in Alzheimer mouse models by targeting mGluR5 to prevent synaptic tagging by C1Q. Sci Transl Med.14(647):eabi8593.

66. Sun W, McConnell E, Pare JF, Xu Q, Chen M, Peng W, Lovatt D, Han X, Smith Y, Nedergaard M (2013) Glutamate-dependent neuroglial calcium signaling differs between young and adult brain. Science.339(6116):197–200.

67. Swanson CJ, Bures M, Johnson MP, Linden AM, Monn JA, Schoepp DD (2005) Metabotropic glutamate receptors as novel targets for anxiety and stress disorders. Nat Rev Drug Discov.4(2):131–44.

68. Tanzi RE (2005) The synaptic Abeta hypothesis of Alzheimer disease. In: Nat Neurosci, pp. 977–9: United States.

69. Tewes B, Frehland B, Schepmann D, Schmidtke KU, Winckler T, Wünsch B (2010) Design, Synthesis, and Biological Evaluation of 3-Benzazepin-1-ols as NR2B-Selective NMDA Receptor Antagonists. ChemMedChem.5(5):687–95.

70. Treyer V, Gietl AF, Suliman H, Gruber E, Meyer R, Buchmann A, Johayem A, Unschuld PG, Nitsch RM, Buck A, Ametamey SM, Hock C (2020) Reduced uptake of [11C]-ABP688, a PET tracer for metabolic glutamate receptor 5 in hippocampus and amygdala in Alzheimer’s dementia. Brain Behav.10(6):e01632.

71. Tu JC, Xiao B, Naisbitt S, Yuan JP, Petralia RS, Brakeman P, Doan A, Aakalu VK, Lanahan AA, Sheng M, Worley PF (1999) Coupling of mGluR/Homer and PSD-95 complexes by the Shank family of postsynaptic density proteins. Neuron.23(3):583–92.

72. Tzioras M, McGeachan RI, Durrant CS, Spires-Jones TL (2023) Synaptic degeneration in Alzheimer disease. Nat Rev Neurol.19(1):19–38.

73. Um Ji W, Kaufman Adam C, Kostylev M, Heiss Jacqueline K, Stagi M, Takahashi H, Kerrisk Meghan E, Vortmeyer A, Wisniewski T, Koleske Anthony J, Gunther Erik C, Nygaard Haakon B, Strittmatter Stephen M (2013) Metabotropic Glutamate Receptor 5 Is a Coreceptor for Alzheimer Aβ Oligomer Bound to Cellular Prion Protein. Neuron.79(5):887–902.

74. Varlow C, Murrell E, Holland JP, Kassenbrock A, Shannon W, Liang SH, Vasdev N, Stephenson NA (2020) Revisiting the Radiosynthesis of [(18)F]FPEB and Preliminary PET Imaging in a Mouse Model of Alzheimer’s Disease. Molecules.25(4).

75. Vizcarra EA, Ulu A, Landrith TA, Qiu X, Godzik A, Wilson EH (2023) Group 1 metabotropic glutamate receptor expression defines a T cell memory population during chronic Toxoplasma infection that enhances IFN-gamma and perforin production in the CNS. Brain Behav Immun.114:131–43.

76. Wang J, He Y, Chen X, Huang L, Li J, You Z, Huang Q, Ren S, He K, Schibli R, Mu L, Guan Y, Guo Q, Zhao J, Xie F (2024) Metabotropic glutamate receptor 5 (mGluR5) is associated with neurodegeneration and amyloid deposition in Alzheimer’s disease: A [(18)F]PSS232 PET/MRI study. Alzheimers Res Ther.16(1):9.

77. Wang J, Huang Q, He K, Li J, Guo T, Yang Y, Lin Z, Li S, Vanderlinden G, Huang Y, Van Laere K, Guan Y, Guo Q, Ni R, Li B, Xie F (2024) Presynaptic density determined by SV2A PET is closely associated with postsynaptic metabotropic glutamate receptor 5 availability and independent of amyloid pathology in early cognitive impairment. Alzheimers Dement.

78. Warnock G, Sommerauer M, Mu L, Pla Gonzalez G, Geistlich S, Treyer V, Schibli R, Buck A, Krämer SD, Ametamey SM (2018) A first-in-man PET study of [(18)F]PSS232, a fluorinated ABP688 derivative for imaging metabotropic glutamate receptor subtype 5. Eur J Nucl Med Mol Imaging.45(6):1041–51.

79. Williams JB, Cao Q, Yan Z (2021) Transcriptomic analysis of human brains with Alzheimer’s disease reveals the altered expression of synaptic genes linked to cognitive deficits. Brain Commun.3(3):fcab123.

80. Wong DF, Waterhouse R, Kuwabara H, Kim J, Brašić JR, Chamroonrat W, Stabins M, Holt DP, Dannals RF, Hamill TG, Mozley PD (2013) 18F-FPEB, a PET radiopharmaceutical for quantifying metabotropic glutamate 5 receptors: a first-in-human study of radiochemical safety, biokinetics, and radiation dosimetry. J Nucl Med.54(3):388–96.

81. Zhang Y, Chen K, Sloan SA, Bennett ML, Scholze AR, O’Keeffe S, Phatnani HP, Guarnieri P, Caneda C, Ruderisch N, Deng S, Liddelow SA, Zhang C, Daneman R, Maniatis T, Barres BA, Wu JQ (2014) An RNA-sequencing transcriptome and splicing database of glia, neurons, and vascular cells of the cerebral cortex. J Neurosci.34(36):11929–47.

82. Zhang Y, Sloan SA, Clarke LE, Caneda C, Plaza CA, Blumenthal PD, Vogel H, Steinberg GK, Edwards MS, Li G, Duncan JA, 3rd, Cheshier SH, Shuer LM, Chang EF, Grant GA, Gephart MG, Barres BA (2016) Purification and Characterization of Progenitor and Mature Human Astrocytes Reveals Transcriptional and Functional Differences with Mouse. Neuron.89(1):37–53.

83. Zhang Y, Sloan SA, Clarke LE, Caneda C, Plaza CA, Blumenthal PD, Vogel H, Steinberg GK, Edwards MSB, Li G, Duncan JA, 3rd, Cheshier SH, Shuer LM, Chang EF, Grant GA, Gephart MGH, Barres BA (2016) Purification and Characterization of Progenitor and Mature Human Astrocytes Reveals Transcriptional and Functional Differences with Mouse. Neuron.89(1):37–53.

84. Zhang YN, Fan JK, Gu L, Yang HM, Zhan SQ, Zhang H (2021) Metabotropic glutamate receptor 5 inhibits α-synuclein-induced microglia inflammation to protect from neurotoxicity in Parkinson’s disease. J Neuroinflammation.18(1):23.

