## Supplemental files for "Hippocampal mGluR5 levels are comparable in Alzheimer’s and control brains, and divergently influenced by amyloid and tau in control brain"

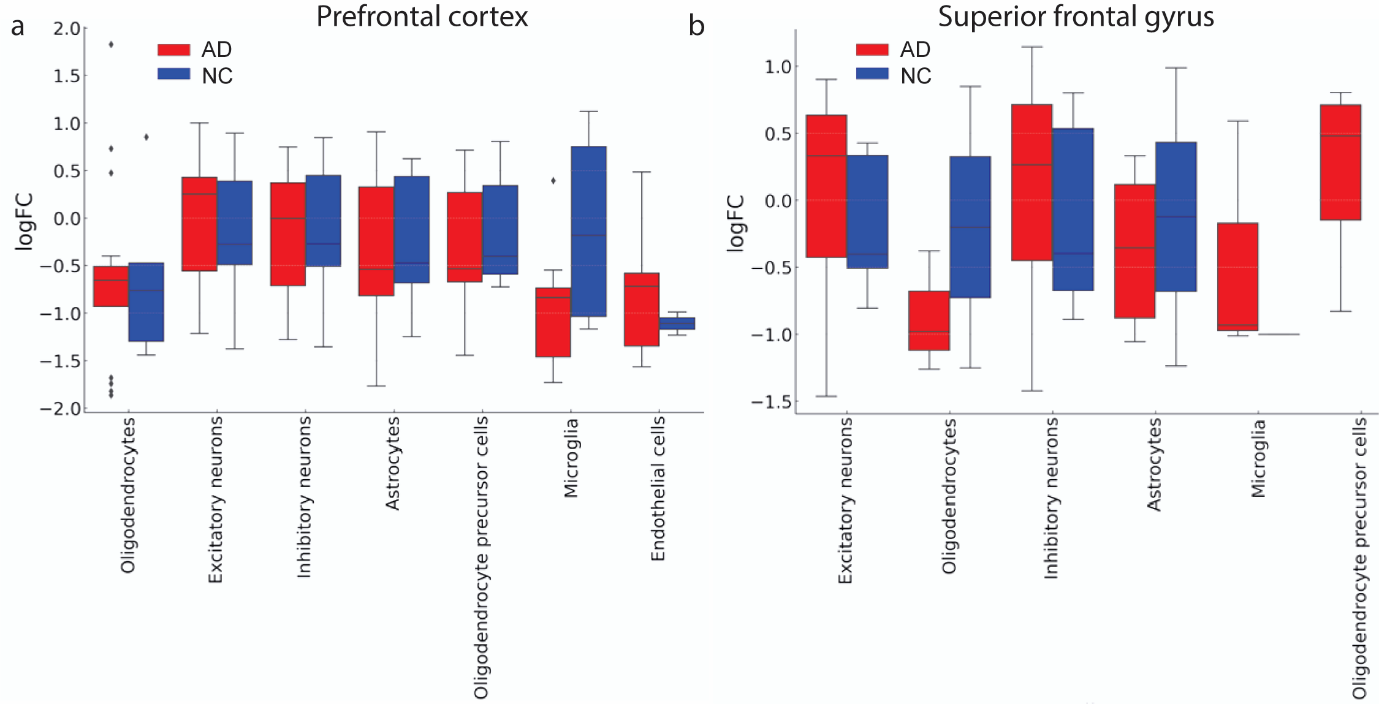

**SFig. 1**: **GRM5 RNA levels in the prefrontal cortex and superior frontal gyrus (BA) of the AD and NC groups**. (**a**) Oligodendrocytes (U=41.0000, p=0.9396, AD n=17, NC n=5), excitatory neurons (U=2215.0000, p=0.7984, AD n=83, NC n=52), inhibitory neurons (U=1070.0000, p=0.6739, AD n=49, NC n=46), astrocytes (U=93.0000, p=0.4957, AD n=20, NC n=11), oligodendrocyte precursor cells (U=124.0000, p=0.0422, AD n=21, NC n=19), microglia (U=10.0000, p=0.2601, AD n=9, NC n=4), and endothelial cells (U=10.0000, p=0.7111, AD n=8, NC n=2) (**b**) in the superior frontal gyrus of the AD and NC; oligodendrocytes (U=2.0000, p=0.8000, AD n=3, NC n=2), excitatory neurons (U=339.0000, p=0.1369, AD n=44, NC n=2) logFC: log fold change. Boxplots showing the medians, quartiles, and outliers. Data from the scREAD database (29) (https://bmbls.bmi.osumc.edu/scread/), which compiles comprehensive datasets from GEO (Gene Expression Omnibus) (17) and Synapse.

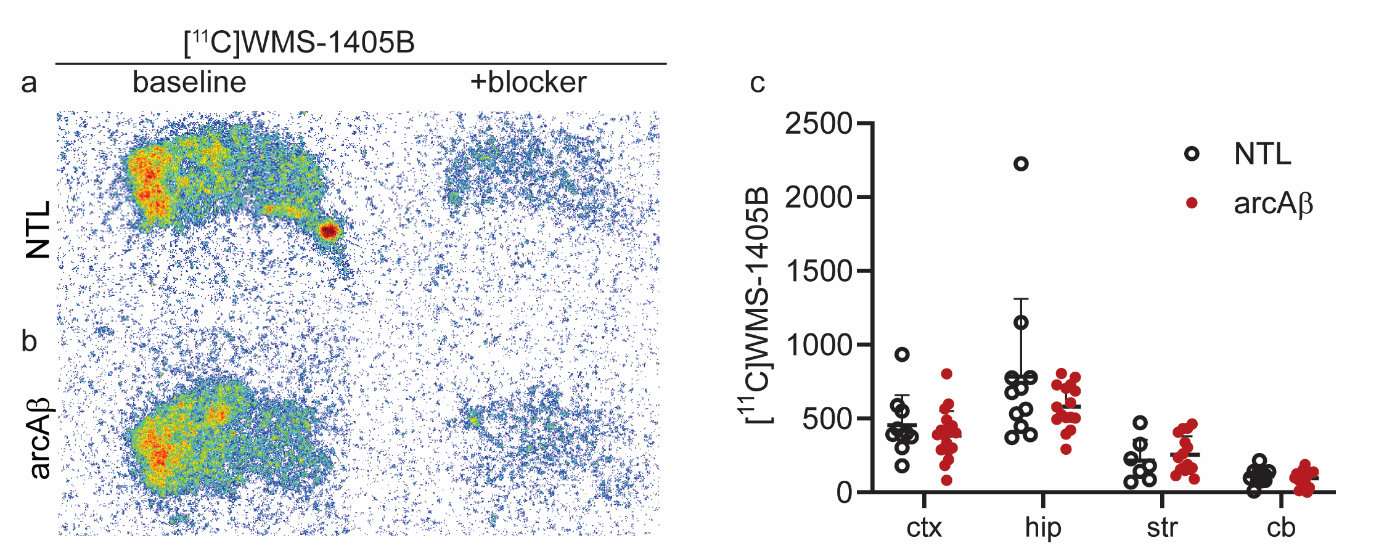

**SFig. 2 Autoradiography of GluN2B expression levels in the brains of arcA**β **mice.** (**a, b**) Representative [^11^C]WMS1405B of GluN2B autoradiographic images of sagittal brain sections of nontransgenic littermates (NTL) and arcAβ mice (baseline and in the presence of blocker). (**c**) Quantification of regional [^11^C]WMS1405B binding in the cortex (ctx), hippocampus (hip), striatum (str), and cerebellum (cb). Nonparametric Mann‒Whitney test, arcAβ vs. NTL.

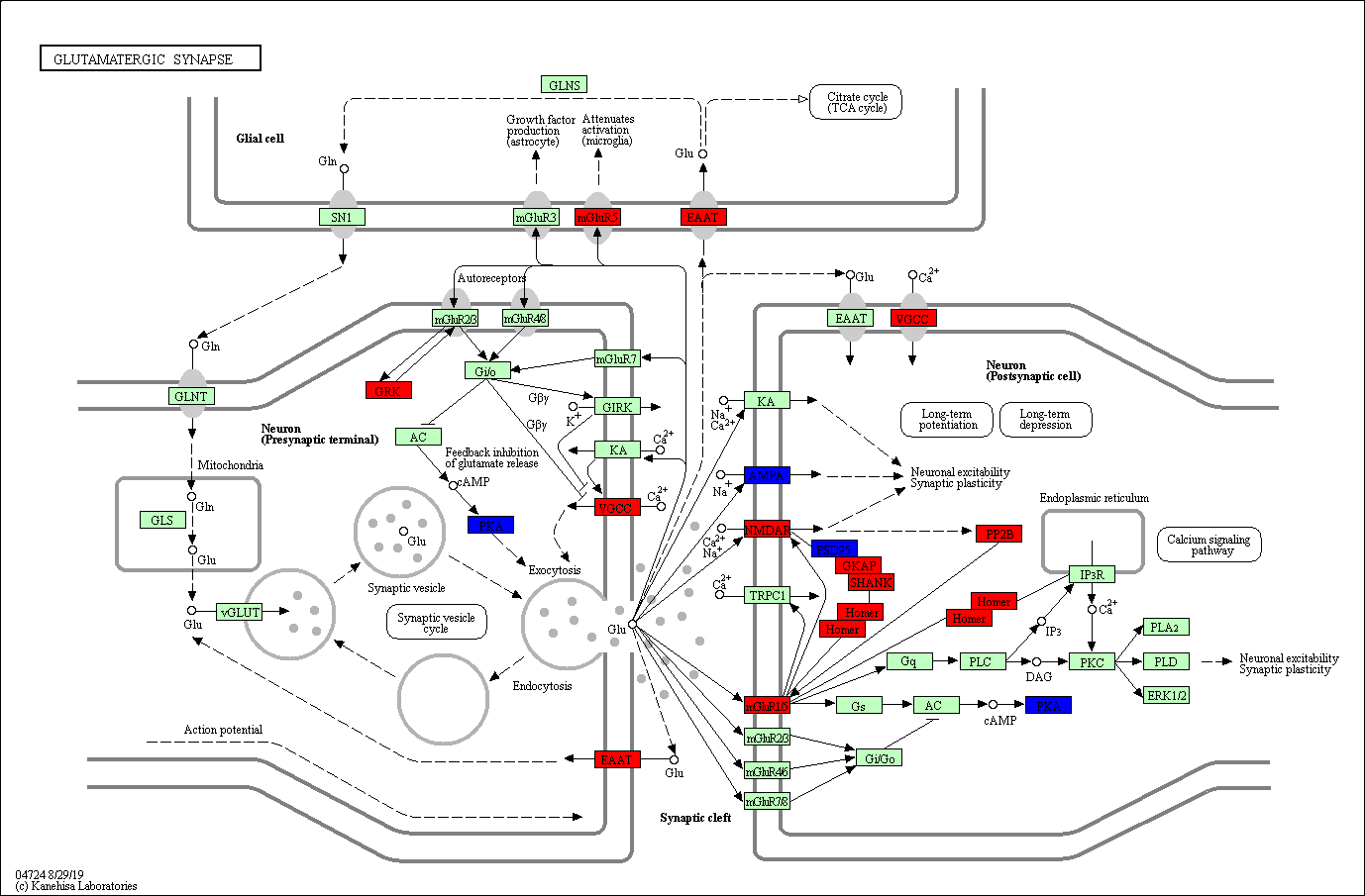

**SFig. 3 Changes in the glutamatergic synapse pathway in the 3×Tg mouse brain according to the proteomics results.**

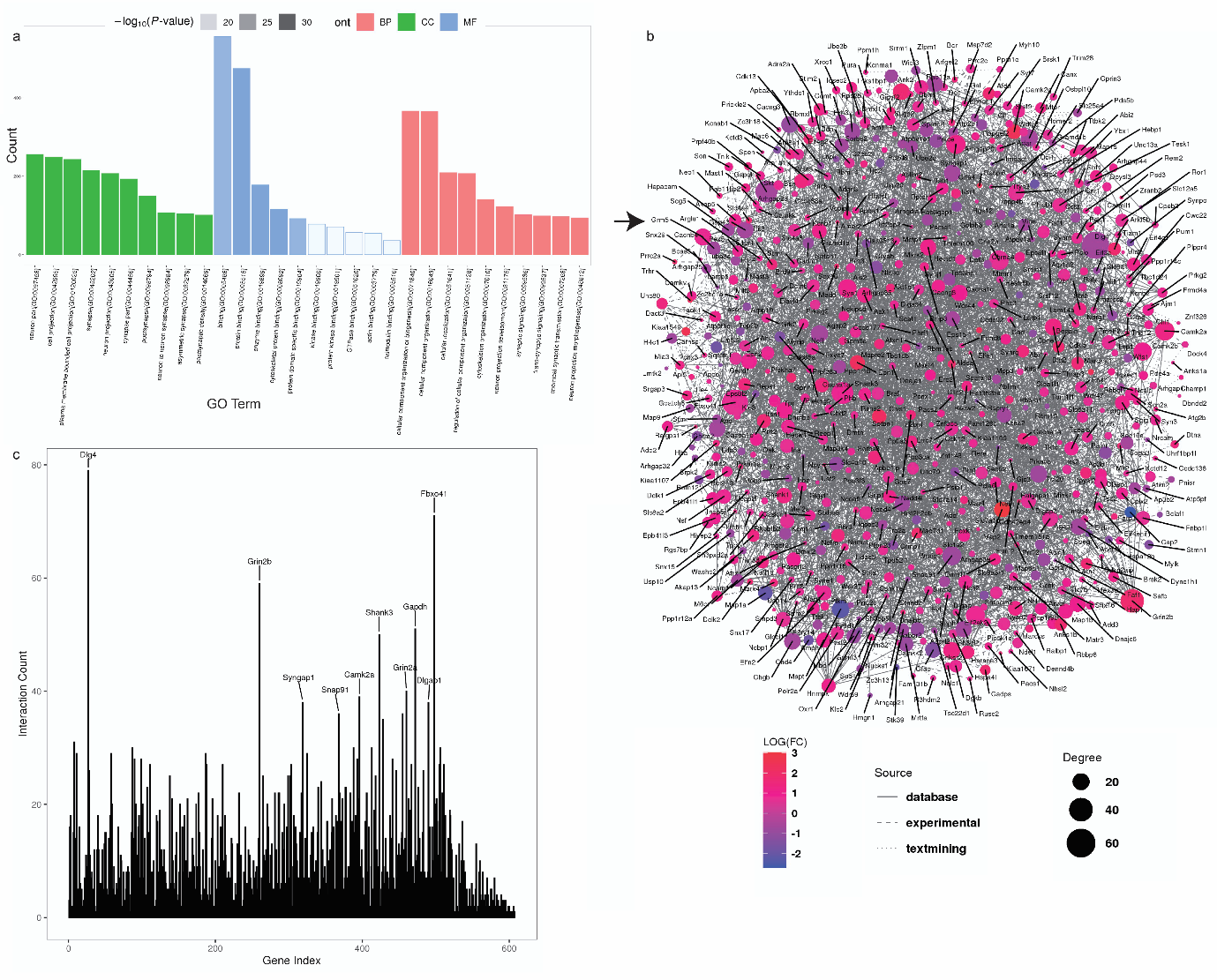

**SFig. 4 Protein–protein interactions and Gene Ontology enrichment analysis in the hippocampi of 16-month-old 3×Tg and WT mice**. (a) Gene Ontology (GO) enrichment analysis of biological process (BP), cellular component (CC), and molecular function (MF) terms. The x-axis indicates the enrichment count of DEPs. (b, c) Network and histogram analysis of protein–protein interactions.

**STable 1 Summary of the antibodies and chemicals used for staining**

| Item | Catalog no | Dilution | Supplier |
| --- | --- | --- | --- |
| Mouse phospho-Tau (Ser202, Thr205) monoclonal antibody (AT-8, IHC, IF) | MN1020 | 1:1000 | Invitrogen |
| Mouse purified anti-β-Amyloid, 1-16 monoclonal antibody (6E10, IF) | 803001 | 1:1000 | Biolegend |
| Mouse purified anti-β-Amyloid, 17-24 monoclonal antibody (4G8, IHC) | 800701 | 1:4000 | Biolegend |
| Rabbit ionized calcium-binding adapter molecule 1 (Iba1), IHC & IF | 019-19741 | 1:1000 | WAKO |
| Guinea pig GFAP polyclonal antibody | BP5082 | 1:1000 | OriGene |
| Goat purinergic P2X7 receptor (P2X7R) | NBP1-37775 | 1:100 | Novus Biologicals |
| Rabbit Iba-1 polyclonal (IHC) | 019-19741 | 1:1000 | WAKO |
| Rabbit recombinant anti-mGluR5 antibody [EPR2425Y] (ab76316) (IF mouse brain) | ab76316 | 1:1000 | Abcam |
| Recombinant HRP anti-mGluR5 antibody [EPR2425Y] (IF human brain) | ab196482 | 1:100 | Abcam |
| Alexa fluor488 donkey anti-rabbit IgG (H+L) | 711-545-152 | 1:250 | Jackson |
| Alexa fluor488 donkey anti-mouse IgG (H+L) | 715-545-151 | 1:500 | Jackson |
| Donkey anti-Rabbit IgG (H+L) Highly Cross-Adsorbed Secondary Antibody, Alexa Fluor™ 647 |  |  |  |
| Alexa fluor647 donkey anti-mouse IgG (H+L) | 715-605-151 | 1:250 | Jackson |
| DAPI (4',6-diamidino-2-Phenylindole, Dihydrochloride) | D1306 | 1:1000 | Invitrogen |

**STable 2 Materials used in the proteomic analysis**

| Item | Comapny |
| --- | --- |
| Urea, analytically pure | GibcoBRL |
| Sequencing grade Trypsin | Promega |
| BCA protein quantification kit | Fisher Scientific |
| Trifluoroacetic acid (TFA) | Sigma |
| Ammonium formate | Sigma |
| TMT6 labeling kit | Fisher Scientific |
| PMSF, ultra pure grade | Amesco |
| Ethylenediaminetetraacetic acid (EDTA), ultra pure grade | Amesco |
| 4-hydroxyethylpiperazineethanesulfonic acid (HEPES) | Sigma |
| Protease inhibitors | Roche, Switzerland |
| Dithiothreitol (DTT), chemically pure | Promega |
| Iodoacetamide (IAA), chemically pure | Promega |
| Sodium dodecyl sulfate, chemically pure | Sigma |
| Ethanol, mass spectrometry pure | Fisher Scientific |
| Formic acid, mass spectrometry pure | Fisher Scientific |
| Acetonitrile, mass spectrometry pure | Fisher Scientific |
| Acetone, mass spectrometry pure | Fisher Scientific |
| Triethylammonium bicarbonate (TEAB) | Santa Cruz |
| 25% Ammonia | Santa Cruz |
| sep-Pak C18 desalting column, 1 cc (100 mg) | Waters Corporation |
| High-Select™ Fe-NTA Phosphopeptide Enrichment Kit, Catalog Number A32992 | Fisher Scientific |
| High pH reversed-phase column, Acquity UPLC®BEH C18 1.7 µm, 2.1×50 mm | Waters Corp. |
| Nanoscale peptide analysis column, Acclaim PepMap C18, 75 µm×250 mm | Thermo Scientific |

**STable 3: The loadings of the first principal component for the AD and NC datasets.**

| Dataset | Variable | Loading |
| --- | --- | --- |
| AD | tau_CA1 | 0.324831582 |
| AD | tau_CA2/3 | 0.3244096 |
| AD | tau_DG | 0.320446357 |
| AD | tau_SUB | 0.336732901 |
| AD | tau_EC | 0.333922802 |
| AD | mGluR5_CA1 | 0.297966626 |
| AD | mGluR5_CA2/3 | 0.303314641 |
| AD | mGluR5_DG | 0.312250017 |
| AD | mGluR5_SUB | 0.298191548 |
| AD | mGluR5_EC | 0.307345651 |
| NC | tau_CA1 | 0.295721449 |
| NC | tau_CA2/3 | 0.30396916 |
| NC | tau_DG | 0.297296995 |
| NC | tau_SUB | 0.298753634 |
| NC | tau_EC | 0.321782801 |
| NC | mGluR5_CA1 | -0.330733374 |
| NC | mGluR5_CA2/3 | -0.327364085 |
| NC | mGluR5_DG | -0.333040807 |
| NC | mGluR5_SUB | -0.331224506 |
| NC | mGluR5_EC | -0.319046243 |

**STable 4: The loadings of the first canonical variables for the AD and NC datasets.**

| Region | Loading | Type |
| --- | --- | --- |
| mGluR5_CA1 | -0.026997499 | AD_mGluR5 |
| mGluR5_CA2/3 | -0.026626566 | AD_mGluR5 |
| mGluR5_DG | -0.045730013 | AD_mGluR5 |
| mGluR5_SUB | 0.019290684 | AD_mGluR5 |
| mGluR5_EC | 0.067981157 | AD_mGluR5 |
| tau_CA1 | 0.006089536 | AD_tau |
| tau_CA2/3 | -0.075591698 | AD_tau |
| tau_DG | -0.030875443 | AD_tau |
| tau_SUB | 0.033022642 | AD_tau |
| tau_EC | 0.060984105 | AD_tau |
| mGluR5_CA1 | -0.136708071 | NC_mGluR5 |
| mGluR5_CA2/3 | -0.010637764 | NC_mGluR5 |
| mGluR5_DG | 0.063482794 | NC_mGluR5 |
| mGluR5_SUB | 0.070333901 | NC_mGluR5 |
| mGluR5_EC | 0.00304709 | NC_mGluR5 |
| tau_CA1 | -0.128040406 | NC_tau |
| tau_CA2/3 | -0.006962965 | NC_tau |
| tau_DG | 0.039084885 | NC_tau |
| tau_SUB | 0.032048738 | NC_tau |
| tau_EC | 0.052963155 | NC_tau |
